# mSWI/SNF promotes polycomb repression both directly and through genome-wide redistribution

**DOI:** 10.1101/2020.01.29.925586

**Authors:** Christopher M. Weber, Antonina Hafner, Simon M. G. Braun, Jacob G. Kirkland, Benjamin Z. Stanton, Alistair N. Boettiger, Gerald R. Crabtree

**Affiliations:** Department of Pathology, Stanford University School of Medicine, Stanford, California, USA; Developmental Biology, Stanford University School of Medicine, Stanford, California, USA; Nationwide Children’s Hospital, Center for Childhood Cancer and Blood Diseases, Columbus, Ohio, USA; Department of Pediatrics, The Ohio State University College of Medicine, Columbus, Ohio, USA; Department of Biological Chemistry and Pharmacology, The Ohio State University College of Medicine, Columbus, Ohio, USA; Howard Hughes Medical Institute, Chevy Chase, Maryland, USA

**Keywords:** transcriptional regulation, ATP-dependent chromatin remodeling, SWI/SNF complex, polycomb, PRC1, PRC2, *Hox* genes, targeted protein degradation, degron

## Abstract

The mammalian SWI/SNF, or BAF complex, has a conserved and direct role in antagonizing polycomb-mediated repression. Yet, BAF appears to also promote repression by polycomb in stem cells and cancer. How BAF both antagonizes and promotes polycomb-mediated repression remains unknown. Here, we utilize targeted protein degradation to dissect the BAF-polycomb axis in embryonic stem cells on the timescale of hours. We report that rapid BAF depletion redistributes both PRC1 and PRC2 complexes from highly occupied domains, like *Hox* clusters, to weakly occupied sites that are normally opposed by BAF. Polycomb redistribution from highly repressed domains results in their decompaction, gain of active epigenomic features, and transcriptional derepression. Surprisingly, through dose-dependent degradation of PRC1 & PRC2 we identify both a conventional role for BAF in polycomb-mediated repression and a second mechanism acting by global redistribution of polycomb. These findings provide new mechanistic insight into the highly dynamic state of the Polycomb-Trithorax axis.

## Introduction

Chromatin regulation is critical to establish and maintain the precise gene expression states that define cellular identity and prevent human pathologies (reviewed in ref. ^1^). The balance between different states primarily involves the antagonism between activating (Trithorax-group) and repressive (Polycomb-group) proteins (reviewed in ref. ^2^). Opposition between these two classes of chromatin regulators was first demonstrated genetically during Drosophila development (reviewed in ref. ^3^). For example, deletion of Polycomb-group genes results in *Hox* gene derepression which gives rise to homeotic transformations, whereas Trithorax-group mutations dominantly suppress these phenotypes^4,5^. Similarly, loss of function mutations in Trithorax-group genes also produce homeotic transformations but are instead due to insufficient *Hox* gene expression^6^. Following these pioneering studies, the genes encoding members of these groups have been shown to be mutated in many human diseases. This is especially true for the BAF (mSWI/SNF) complex, a Trithorax-group homolog, which is frequently mutated in many cancers, neurodevelopmental disorders, and intellectual disabilities^7–9^.

BAF complexes are combinatorially assembled chromatin remodeling enzymes of ~15 subunits that hydrolyze ATP to mobilize nucleosomes and generate accessible DNA^10^. In mammalian cells, BAF directly evicts polycomb repressive complex 1 (PRC1)^11,12^ leading to transcriptional derepression^13^. Thus, the ability of Trithorax-group proteins to antagonize polycomb-mediated repression is conserved from Drosophila to mammals. PRC1 and PRC2 direct H2A ubiquitination (H2AK119ub1) and trimethylation of histone H3 at lysine 27 (H3K27me3) respectively and spatially constrain the genome in support of transcriptional repression (reviewed in ref. ^2^). BAF-mediated polycomb antagonism is essential during development and is thought to underlie the tumor suppressive role in human cancers (reviewed in ref. ^14^).

Conversely, BAF’s potent ability to antagonize polycomb-mediated repression is coopted in synovial sarcoma, where fusion of SSX onto the SS18 subunit retargets BAF and opposes polycomb-mediated repression to drive tumor growth^7,15^.

Despite extensive evidence that BAF has a dominant role in polycomb opposition, BAF appears to also be required for polycomb-mediated repression in embryonic stem cells, during lineage commitment, and in sub-types of rare atypical teratoid rhabdoid tumors (ATRT) that are characterized by *Hox* gene derepression^16–18^. ATRTs are highly malignant tumors that are typically seen in children younger than 3. These tumors are characterized by inactivation of either *SMARCB1* (*BAF47*) or *SMARCA4* (*BRG1*), the core ATPase subunit, and rarely contain mutations in other genes^19,20^. Currently, the mechanism by which BAF simultaneously supports active and repressed states remains unclear.

A major limitation to resolving this question has been with the use of loss of function approaches that lack sufficient temporal resolution to distinguish primary from secondary effects. Chromatin regulators tend to be stable for several days following conventional depletion methods and are subject to numerous feedback mechanisms. This is especially problematic when studying chromatin remodelers like BAF, which regulate accessibility for many DNA-based processes and have numerous ascribed roles from transcriptional regulation to cell division. To overcome these limitations, we implemented a chemical genetic approach to enable rapid targeted protein degradation of essential BAF, PRC1, and PRC2 subunits in mouse embryonic stem cells (mESCs). We demonstrate that targeted degradation of the BAF ATPase subunit directly derepresses many genes that are highly occupied by polycomb, such as *Hox* genes and developmental regulators, that is immediately coincident with depletion kinetics. We show that BAF depletion results in the genome-wide redistribution of PRC1 & PRC2, independent of transcription, from highly occupied domains like *Hox* clusters to sites that are normally opposed by BAF, resulting in their physical decompaction. Through dose-dependent degradation of PRC1 & PRC2 we also identify a direct role for BAF in facilitating polycomb-mediated repression. Our mechanistic study reconciles the dual role for BAF in opposition and maintenance of polycomb-mediated repression, highlighting the dynamic nature of the Polycomb-Trithorax axis with implications for human disease.

## Results

### Brg1 degradation with auxin is rapid and near complete

To temporally resolve the effects of BAF inactivation, we developed an ESC line where the sole ATPase subunit Brg1 (also known as Smarca4) can be rapidly degraded. Both endogenous alleles of *Brg1* were tagged with the minimal 44 amino acid auxin inducible degradation tag (AID*) using CRISPR-Cas9 with homology-dependent repair^21,22^ (Supplementary Fig. 1a,b). The F-box protein osTIR1 was inducibly expressed in these cells, which forms a hybrid SCF ubiquitin ligase complex that targets Brg1 for degradation by the proteasome when the small molecule auxin is added (Fig. 1a). Tagging Brg1 with AID* did not affect protein abundance, such that for all experiments Brg1 levels were equivalent to WT before adding auxin (Supplementary Fig. 1c). Additionally, the cells divided at the same rate and were indistinguishable from the parent mESC line. Consistent with other studies, auxin treatment in the absence of osTIR1 was innocuous to cell viability and growth^23^. Yet, when auxin was added to cells expressing osTIR1, Brg1 was rapidly degraded, with a protein half-life of ~30 minutes and maximal, near-complete, degradation by 2 hours (Fig. 1b,c). This degron strategy resulted in much faster loss of function than genetic deletion (~2h vs. 3 days, Supplementary Fig. 1d) and induced changes to colony morphology at the 24h time-point (Supplementary Fig. 1d,e) so we conducted all subsequent experiments at short time-points (≤ 8 hours); shorter than one cell cycle. Thus, the Brg1 degron is a tractable and robust strategy to resolve the direct effects of BAF inactivation on the timescale of hours.

**Fig. 1:**
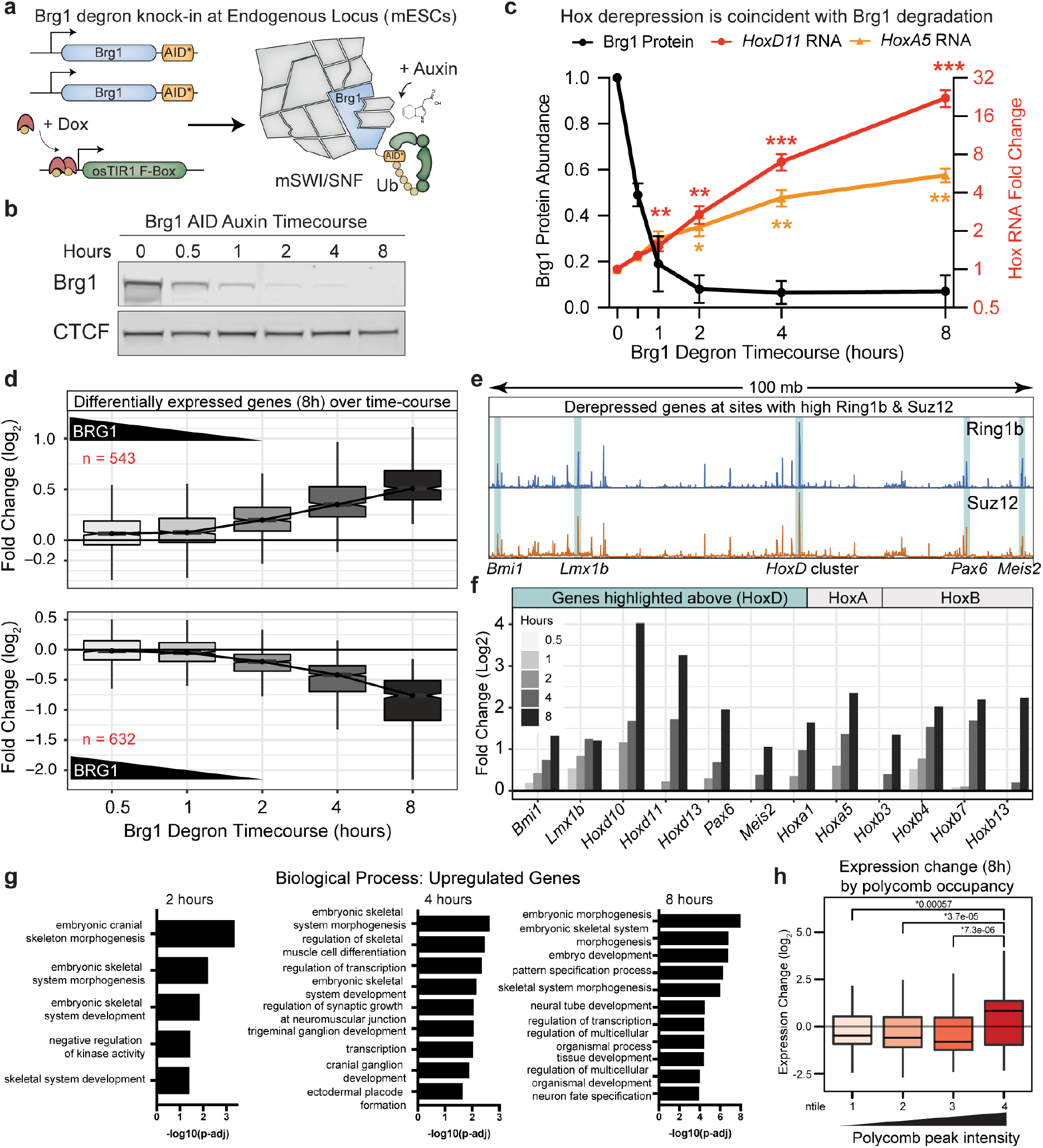
Brg1 degradation results in the derepression of many highly polycomb-bound genes. (a) Schematic depicting Brg1 auxin inducible degradation (AID) strategy in mESCs. Both alleles of Brg1 are endogenously tagged at the C-terminus with the minimal AID* tag. osTIR1 is induced with doxycycline before degrading Brg1 with auxin. (b) Representative western blot showing Brg1 AID degradation kinetics and efficiency. (c) Quantitation of Brg1 degradation kinetics (left axis, Maximal by 2h with T1/2 = ~30m) and induction of *HoxD11* and *HoxA5* transcription before Brg1 is maximally degraded (right axis, fold change by ddCt). Error bars represent mean +/- standard error of the mean from n=3 and n=4 biological replicates, respectively. Significance was calculated by individual Student’s t test (*P ≤ 0.05, **P ≤ 0.01, ***P ≤ 0.001). (d) Fold change in expression level (Log2) for genes that are differentially expressed at 8h (FDR-corrected P < 0.05) across all time-points (increased n=543, decreased n=632) with Brg1 degradation kinetics overlaid. (e) Browser tracks showing Ring1b and Suz12 ChIP-seq signal (0 hour) at 100mb of 181mb chromosome 2. Regions with high polycomb occupancy that become derepressed by Brg1 degradation are highlighted. (f) Timecourse of Log_2_ fold changes for genes highlighted in (g) and genes from *HoxA* and *B* clusters that are derepressed by Brg1 degradation. (h) Gene set enrichment analysis for differentially upregulated genes at 2, 4, and 8 hour timepoints. (h) Genes with high Ring1b and Suz12 occupancy at transcription start site (+/- 2kb) become derepressed by Brg1 degradation (by ntile of peak intensity, Wilcoxon rank-sum p-value is shown). All experiments were conducted in serum/LIF.

### Brg1 degradation results in the derepression of many highly polycomb bound genes

A previous study reported derepression of *Hox* genes following 3 days of Brg1 conditional knockout^18^. This time-point is unable to capture primary effects and undoubtedly combines secondary changes driven by a smaller number of primary events (Supplementary Fig. 2d). We first leveraged the fast Brg1 depletion kinetics to test whether this effect is causal. Indeed, we observed a time-dependent increase in *HoxA5* and *HoxD11* transcription that was directly coincident with Brg1 degradation, visible as early as 0.5h, with both genes becoming significantly derepressed by the time that Brg1 was maximally degraded and expression continued to increase for 8h (*P* < 0.05) (Fig. 1c). In contrast, *Hox* gene derepression was not caused by auxin induced degradation of *Pbrm1*, a defining member of the pBAF complex, controlling for artifacts of the experimental approach (Supplementary Fig. 2b) and also demonstrating that this repressive role is specific to the canonical BAF complex.

To comprehensively define the transcriptional kinetics, we next conducted an RNA-seq timecourse at 0.5, 1, 2, 4, and 8h after degrading Brg1 with auxin. The number of differentially expressed genes increased at each successive time-point with n = 543 upregulated and n = 632 downregulated genes at 8-hours (FDR-corrected *P* < 0.05) (Supplementary Fig. 2a). Consistent with the trend for *HoxA5* and *HoxD11*, the genome-wide response was also directly coincident with Brg1 degradation (Fig. 1d). For genes upregulated at 8h, median log_2_ fold changes were higher as early as 0.5h (~50% Brg1 levels) and increased at each successive time-point. Downregulated genes exhibited a minor time-lag relative to upregulated genes, as expected, considering the time needed for transcript decay and completion of elongation, yet a reduction in transcription was clearly seen by 2h and transcription of these genes continued to decrease at each successive time-point (Fig. 1d, bottom). Most transcriptional changes occurred before Brg1 was even completely degraded, at time-scales ~0.5-2h. Considering that protein maturation time and half-life of protein/RNA is generally much longer than this, we conclude that these changes are the direct result of rapid Brg1 degradation and not a secondary effect.

We were intrigued to find that, in addition to Hox, many of the most strongly derepressed genes colocalized with the strongest PRC1 and PRC2 peaks on the chromosome (Fig. 1e). Previous studies have shown that these sites form extra long-range chromosomal interactions that require PRC1, for example between *HoxD* and *Bmi1, Pax6, Meis2*, and *Lmx1b*^24^. *HoxA*, *B*, and *D* genes were amongst the most strongly derepressed and most of these genes showed a time-dependent increase in transcription by ~2h (Fig. 1f), consistent with the results in Fig. 1c and Fig. 1d. *Bmi1* (*PCGF4*) was also derepressed quickly upon Brg1 degradation and is a reliable BAF repressed gene that has been used as a reporter in a high-throughput inhibition screen^25^. Yet, strong ectopic overexpression did not cause *Hox* gene derepression (Supplementary Fig. 2e), providing further support that derepression within polycomb domains is a direct cause of rapid Brg1 depletion.

In ESCs, the polycomb system represses developmental regulators, including many transcription factors, to prevent differentiation^26^. In light of our finding that many polycomb-bound genes were derepressed by Brg1 depletion, we next conducted gene set enrichment analysis (Fig. 1g). As expected, we find that many patterning, differentiation, and developmental categories were enriched, even at the 2h time-point (see Supplementary Fig. 2c for downregulated genes). Considering that many genes with high polycomb occupancy became derepressed, like the *Hox* clusters, we sought to determine if this is a more general trend that can be predicted by polycomb occupancy. At 8h, where 1175 genes are differentially expressed, we found that weakly polycomb bound genes (average Ring1b and Suz12 intensity +/- 2kb from the TSS, ntiles 1-3) tended to become repressed, whereas genes with the highest PRC1 and PRC2 levels (ntile 4) tended to become derepressed by Brg1 degradation (Fig. 1h, *P* < 0.05 between ntile 4 and 1-3). Thus, rapid Brg1 depletion has opposing effects on polycomb-mediated gene regulation, with derepression of highly and repression of lowly polycomb occupied genes.

### PRC1 and PRC2 are quickly redistributed upon Brg1 degradation

We observed quick transcriptional derepression within polycomb domains, so we next sought to determine the effect of Brg1 degradation on PRC1&2 occupancy genome-wide by conducting ChIP-seq for Ring1b and Suz12 (core subunits of PRC1 and PRC2 complexes) (Supplementary Fig. 3a). Consistent with the established role for BAF in evicting polycomb, we found more Ring1b and Suz12 peaks were increased than decreased upon 8h of Brg1 degradation (for Ring1b 931 increased and 641 decreased, where 1,572/10,569 (14.9%) of peaks were differentially bound and for Suz12 457 increased, 200 decreased, where 657/8,853 (7.4%) of peaks were differentially bound at FDR-corrected *P* < 0.1). Differential peaks with low Ring1b and Suz12 occupancy (as assessed by normalized peak counts) were mostly increased and > 90% were bound by BAF, whereas highly occupied peaks mostly decreased following 8h Brg1 degradation (Fig. 2a). This general trend can be seen in representative genome-browser snapshots of increased (*Cyp2s1* and *Ptk2b*) and decreased (*HoxA/D* clusters) sites (Fig. 2b) (Supplementary Fig. 3b,c). Collectively, these results demonstrate that the loss of PRC2 at *Hox* clusters, previously seen after 72h in the Brg1 conditional deletion^18^, occurs coincident with Brg1 removal and revealed that not only is PRC2 lost, but PRC1 was also depleted. In fact, Brg1 degradation induced changes to PRC1 and PRC2 were highly correlated across all peaks (R = 0.79, p < 2.2e-16) and displayed a high degree of overlap between peaks (Fig. 2c,d). Importantly, these changes were independent of global changes to the core proteins and the histone modifications placed by these complexes, even at much longer time-points (Supplementary Fig. 3e).

**Fig. 2:**
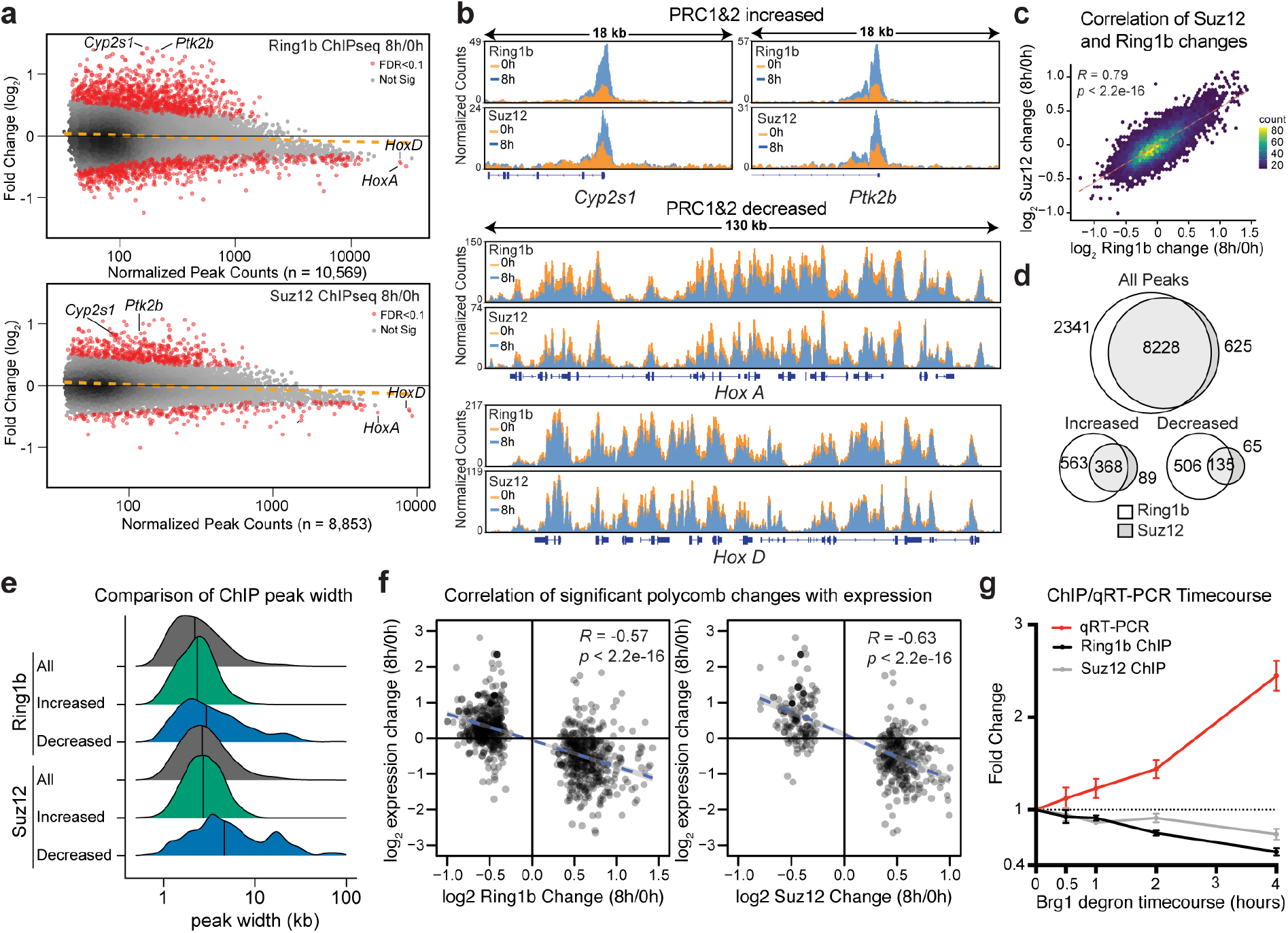
PRC1 and PRC2 are quickly redistributed upon Brg1 degradation. (a) MA plot showing genome-wide changes to Ring1b and Suz12 ChIP-seq peaks (n = 10,569 and n = 8,853 respectively) at 8h Brg1 degradation with differentially bound peaks in red (FDR-corrected P < 0.1) from four biological replicates. For Ring1b, there are n = 930 increased and n = 640 decreased peaks, and for Suz12 there are n = 457 increased and n = 198 decreased peaks. (b) Representative genome-browser tracks for peaks labeled in (a) that increase (Cyp2s1 and Ptk2b) and decrease (*HoxA* and *HoxD* clusters) normalized by read-depth. (c) Correlation plot between Ring1b and Suz12 peak changes as a hexagonal heatmap of 2D bin counts. Correlation and p-values were obtained from Pearson’s product moment correlation. (d) Peak overlap for all, increased, and decreased between Ring1b and Suz12. (e) Peak widths for all, increased, and decreased peak changes for Ring1b and Suz12. (f) Correlation of significantly changed Ring1b and Suz12 peaks that are +/- 2kb from gene transcription start sites and gene expression changes at 8h Brg1 degradation. Correlation and p-values were obtained from Pearson’s product moment correlation. (g) ChIP-qPCR and qRT-PCR timecourse at *Bmi1* locus showing coincident PRC1/2 loss and derepression, error bars depict mean +/- standard error of the mean from three biological replicates. All experiments were conducted in serum/LIF.

Chromatin state enrichment analysis^27^ revealed that regions where PRC1&2 decreased upon Brg1 degradation, were highly enriched for bivalent chromatin, with slight enrichment for the fully repressed state (Supplementary Fig. 3g). We wondered whether the high normalized peak counts seen at decreased sites were due to enrichment for broader domains as a general principle, as seen at the *Hox* clusters, so we plotted the peak width density for all, increased, and decreased peaks (Fig. 2e). We found that the increased peaks had a very similar distribution to all PRC1 and PRC2 peaks (Ring1b median for all peaks = 2.2kb, increased = 2.3kb and Suz12 median for all peaks = 2.9kb, increased = 2.7kb), but decreased peaks initially contained broader polycomb domains (Ring1b median decreased = 2.9kb, Suz12 median decreased = 4.7kb). We previously showed that the basal occupancy level of PRC1 and PRC2 was predictive of the direction of differential gene expression when Brg1 was degraded (Fig. 1h). We next sought to determine how global polycomb changes influence transcription at the same 8h time-point. We found that for all genes, there was a modest but highly significant negative correlation between changes to polycomb occupancy and transcription (Supplementary Fig. 3f, Ring1b: *R* = −0.31, *P* < 2.2e-16, Suz12: *R*= −0.33, *P* < 2.2e-16). Yet, considering only differential peaks, we observed an even stronger negative correlation (*R* = −0.57, *P* < 2.2e-16 and *R* = −0.63, *P* < 2.2e-16 for Ring1b and Suz12 respectively) (Fig. 2f). These results suggest that polycomb redistribution leads to transcriptional derepression. To define the temporal dynamics, we conducted ChIP qPCR and qRT-PCR at Bmi1, a canonical BAF and polycomb repressed gene. Here, polycomb loss and transcriptional derepression were coincident, with subtle Ring1b and Suz12 loss, as early as 0.5h where Brg1 levels are ~50%, that continued to decrease in a time-dependent manner, consistent with polycomb loss leading to derepression. (Fig. 2g, see also Supplementary Fig 3d for longer time-courses). Altogether, these results demonstrate that BAF activity is constantly required to evict polycomb from lowly occupied bivalent sites in order to accumulate at highly occupied broad domains, like *Hox* clusters.

### Brg1 degradation and polycomb loss result in decompaction of *Hox*

Polycomb complexes are known to constrain and physically compact chromatin, which is thought to facilitate repression^28–30^, leading us to ask if the partial PRC1 and PRC2 loss from high polycomb occupancy sites is sufficient to spatially decompact them. To test this, we conducted ORCA experiments (Optical Reconstruction of Chromatin Architecture)^28,31,32^ at *HoxA* and *HoxD* clusters, which lose polycomb and become transcriptionally derepressed by Brg1 degradation. ORCA enables reconstruction of chromatin trajectories (100-700kb) by tiling short regions (2-10kb) with unique barcodes and measuring their nanoscale 3D positions (Fig. 3a,b). We tiled 290kb at *HoxA* and 234kb at *HoxD* in 5kb steps, i.e. different barcode for each 5kb step, that completely cover the polycomb repressed domains and extend into the flanking regions (Fig. 3c,d).

**Fig. 3:**
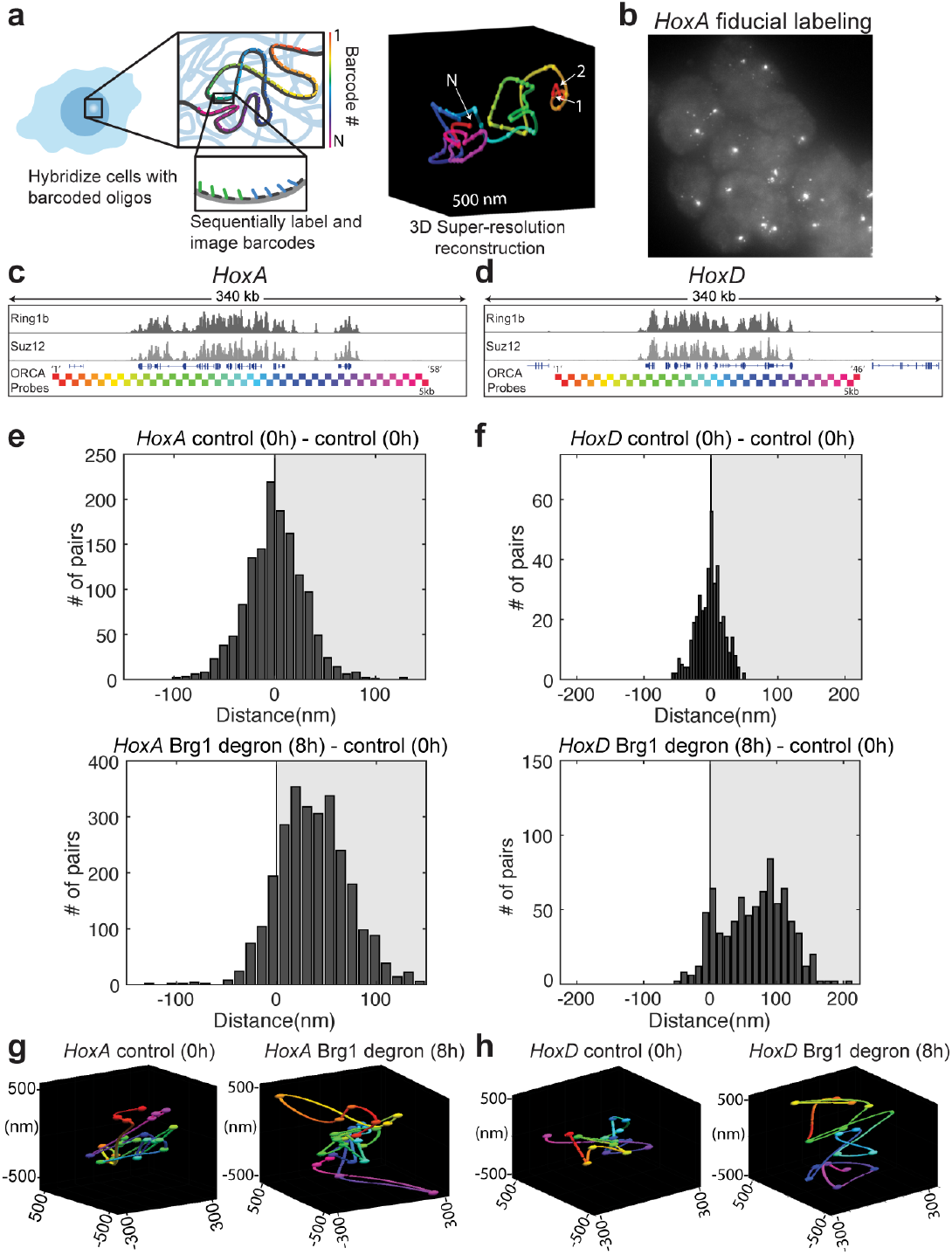
Brg1 degradation and polycomb loss result in decompaction of *HoxA* and *HoxD* regions. (a) Schematic of ORCA method where sequential imaging and labeling of fluorescent barcodes (numbers 1 to N) are artificially color coded by barcode number, enabling reconstruction of the 3D DNA path of a DNA region. (b) mES cells labelled with *HoxA* probe. Each spot corresponds to one allele and shown here is the fiducial staining, that labels the entire 290kb *HoxA* region. c,d) Genomic *HoxA* and *HoxD* regions labelled by ORCA. Bottom tracks illustrate the positions of probes, color coded by barcode number. (e,f) Histograms plot the difference in median distance for all barcode pairs, where median was calculated over all cells, between AID treated and control cells. (g,h) Representative polymers in the control and Brg1 depleted cells for *HoxA* (g) and *HoxD* (h). All experiments were conducted in serum/LIF.

Applying ORCA in the untreated cells and calculating the contact frequency over all cells revealed similar sub-TAD domain architecture and a high correlation with published Hi-C data in mESCs^33^ (Supplementary Fig. 4a,b). We then leveraged the unique ability of ORCA to measure 3D nanoscale distances and quantified the median inter-barcode distance at *HoxA* and *HoxD* following Brg1 degradation. Consistent with redistribution of PRC1 and PRC2 from Hox detected by ChIP, we found higher inter-barcode distances for both *HoxA* and *HoxD* when Brg1 is degraded (8h) (Fig. 3e, f, Supplementary Fig. 4c-e) which is also apparent in the re-constructed polymer traces from single cells (Fig. 3g,h). Thus, the redistribution of polycomb away from *Hox* clusters, despite being incomplete, is sufficient to spatially decompact them.

### Genome-wide polycomb redistribution changes chromatin state from repressed to active

Transcriptional regulation involves a dynamic interplay between activating and repressive forces. We next sought to determine how the genome-wide redistribution of polycomb-mediated repression impacts other chromatin features associated with active transcription. We conducted ChIP-seq for H3K4me3 and H3K27ac following 8h Brg1 degradation. H3K4me3 is a histone modification associated with poised and transcriptionally active genes that is deposited by the COMPASS complex, which has homologous subunits in Drosophila that are Trithorax-group members, like the BAF complex^34^. H3K27ac is primarily deposited at active genes and enhancers by the CBP/p300 complex, which binds to BAF^35^, and antagonizes the mutually exclusive H3K27me3 modification that is placed by PRC2^36^. Interestingly, we found that at 8h Brg1 degradation, both H3K27ac and H3K4me3 were negatively correlated with changes to Ring1b (*R* = −0.45, *P* < 2.2e-16 and *R* = −0.5, *P* < 2.2e-16) and Suz12 (*R* = - 0.37, *P* < 2.2e-16 and *R* = −0.44, *P* < 2.2e-16) genome-wide (Fig. 4a,b), such that where PRC1 and PRC2 increase, active marks were decreased (Fig. 4c) and where PRC1 and PRC2 decreased, active marks increased after Brg1 degradation (Fig. 4d) (Supplementary Fig. 5a,b). These data demonstrate that the global polycomb redistribution induced by Brg1 degradation results in opposite epigenomic changes associated with active transcription.

**Fig. 4:**
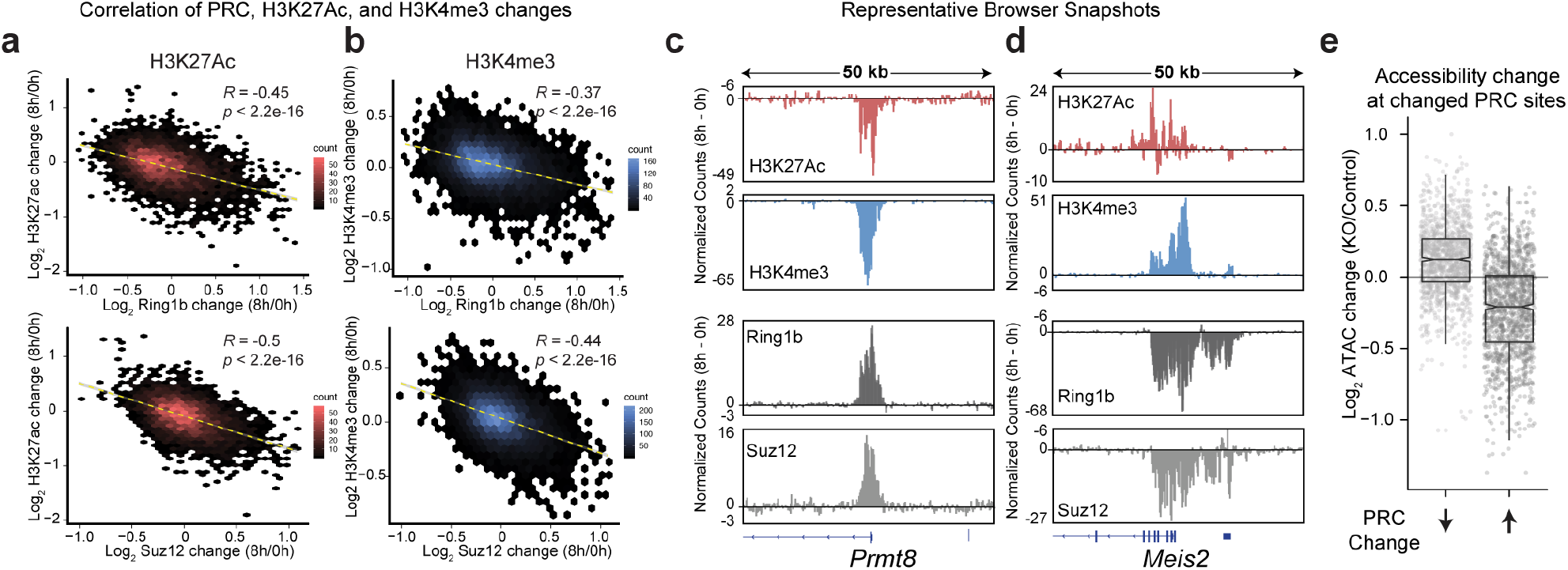
Genome-wide polycomb redistribution changes chromatin state from repressed to active. Correlation plot of fold change (Log2) to H3K27ac (a) H3K4me3 (b) and Ring1b or Suz12 at 8h Brg1 degradation as hexagonal heatmap of 2D bin counts. Correlation and p-values were obtained from Pearson’s product moment correlation. (c) Representative browser snapshots showing *Prmt8* locus that gains Ring1b and Suz12 and loses both H3K27ac and H3K4me3 as the difference in normalized counts (8h - 0h degradation). (d) Browser snapshot at *Meis2* locus which loses Ring1b and Suz12 and gains both H3K27ac and H3K4me3. (e) Boxplot showing the change in accessibility to Tn5 transposition in Brg^fl/fl^ deleted cells (TAM) over control (EtOH), at peaks that have decreased or increased Ring1b or Suz12 signal after 8h Brg1 degradation. (ATAC-seq data from^41^). All experiments were conducted in serum/LIF.

We next sought to understand what drives the global redistribution of polycomb from highly occupied sites. One possibility is that BAF inhibits transcription through nucleosome positioning^37^ which then selectively drives polycomb loss due to physical disruption from the transcription machinery or by competitive removal through preferential binding of PRC2 to RNA^38^. We tested this idea by blocking transcription initiation with Triptolide, +/- auxin, for 8h^39^ (Supplementary Fig. 6a,b). We chose this time-point for consistency with our other ChIP-seq datasets and to minimize toxicity from the inhibitor^40^. We then correlated the changes to Ring1b and Suz12 caused by Brg1 degradation +/- complete inhibition of transcription (auxin + triptolide / triptolide to auxin / DMSO). This experiment revealed that polycomb was still redistributed, despite global transcription inhibition for both Ring1b (*R* = 0.68, *P* < 2.2e-16) and Suz12 (*R* = 0.65, *P* < 2.2e-16) (Supplementary Fig. 6c). Importantly, this correlation was strengthened even further, for peaks that were significantly changed by Brg1 degradation (R = 0.86, *P* < 2.2e-16 and R = 0.83, *P* < 2.2e-16 for Ring1b and Suz12 respectively) (Supplementary Fig. 6d). In contrast, the effect of triptolide vs. auxin alone showed zero, or a very weak correlation (*R* = 0.015 for Ring1b and *R* = 0.25 for Suz12) (Supplementary Fig. 6d). Thus, the polycomb redistribution caused by Brg1 depletion is independent of transcription, consistent with our observation that polycomb is lost across large 100+ kilobase domains, and not just at the genes that become derepressed.

In general, BAF and PRC1 and PRC2 overlap extensively genome-wide, however sites that are highly occupied for either appear mutually antagonistic (Supplementary Fig. 7a). Our polycomb ChIPseq in the Brg1 degron cells showed that PRC1 and PRC2 accumulate at sites that are normally opposed by BAF. To see if the opposite is true, i.e. does PRC1 produce resistance to BAF binding as suggested by prior in vitro experiments^42,43^, we used ES cells in which we could conditionally delete Ring1b^fl/fl^ using a tamoxifen inducible Actin-CreER in a Ring1a^-/-^ background, to completely ablate PRC1 activity^44^. After tamoxifen treatment, we examined the binding of BAF over the ES cell genome by ChIP-seq for the BAF155 core subunit of the BAF complex (Supplementary Fig. 7a,b). In contrast to expectations, we found that while some BAF155 peaks did change, the vast majority >95% were unchanged following PRC1 deletion, and only 1% of differential peaks were within PRC1 domains. (Supplementary Fig. 7c-g). Therefore, the BAF-polycomb axis appears mostly unidirectional, such that BAF antagonizes polycomb at weakly bound sites but is not excluded from highly PRC1 bound sites, like *Hox* clusters.

To investigate this further, we looked at Brg1 dependent accessibility changes by ATAC-seq^41^ at sites where polycomb changes upon loss of Brg1. Accessibility changes reflect SWI/SNF chromatin remodeling activity since the readout is the endpoint of the catalytic cycle, i.e. nucleosome dynamics. We reasoned that if BAF was required to generate accessibility for the DNA-binding subunits that target polycomb complexes, then accessibility changes upon Brg1 loss should mimic polycomb changes. In sharp contrast to this, we found that sites where polycomb was lost became more accessible, and sites that gained polycomb became less accessible (Fig. 4e, and Supplementary Fig. 5c,d). Yet, polycomb doesn’t repress accessibility to Tn5 transposition in mESCs^45,46^, which suggests that Brg1 is required to repress accessibility at these sites. These results are most consistent with a model where BAF facilitates polycomb-mediated repression by repressing accessibility at highly PRC-bound sites, as well as distant PRC1 and PRC2 eviction at abundant sites of low polycomb affinity.

### BAF promotes repression directly and by genome-wide polycomb redistribution

If the BAF driven polycomb redistribution and subsequent decompaction alone is sufficient to drive derepression, we reasoned that this should result in insufficient polycomb to both accumulate and maintain distal repression. To test this, we wanted a system where we could rapidly degrade PRC1&2 in a dose-dependent way and measure the transcriptional response relative to Brg1 depletion at the exact same 8-hour time-point. For this goal, we implemented the dTAG targeted degradation approach, which enables dose-dependent, specific, and efficient protein degradation^47^. We tagged endogenous alleles of essential subunits of PRC1 (Ring1b) and PRC2 (EED) with the FKBP^F36V^ tag that enables targeted degradation in the presence of dTAG13, a heterobifunctional analog of rapamycin. Tagging these proteins didn’t noticeably affect protein abundance, cell viability, or growth (Supplementary Fig. 8a-d).

Treating ESCs with a 10-fold serial dilution of dTAG13 ligand resulted in step-wise, near complete degradation of Ring1b/EED to 75, 50, 12, and 3% of WT levels at the 8h time-point (Fig. 5a) (maximal degradation with a high dTAG13 dose is achieved by 2h, similar to the Brg1 degron). We next conducted RNA-seq from these four polycomb doses at a single 8h time-point, to enable comparison with the Brg1 degron effect. Surprisingly, we found that reducing PRC to 75 and 50% had very modest transcriptional effects. Yet, depleting PRC1&2 to 12% and 3% resulted in many more derepressed genes (n = 179 and n = 970, respectively, FDR-corrected *P* < 0.05) (Fig 5b). Consistent with the statistical cutoff, the magnitude of log2 fold changes also displayed modest dosage sensitivity, until the higher levels of depletion (Fig. 5c).

**Fig. 5:**
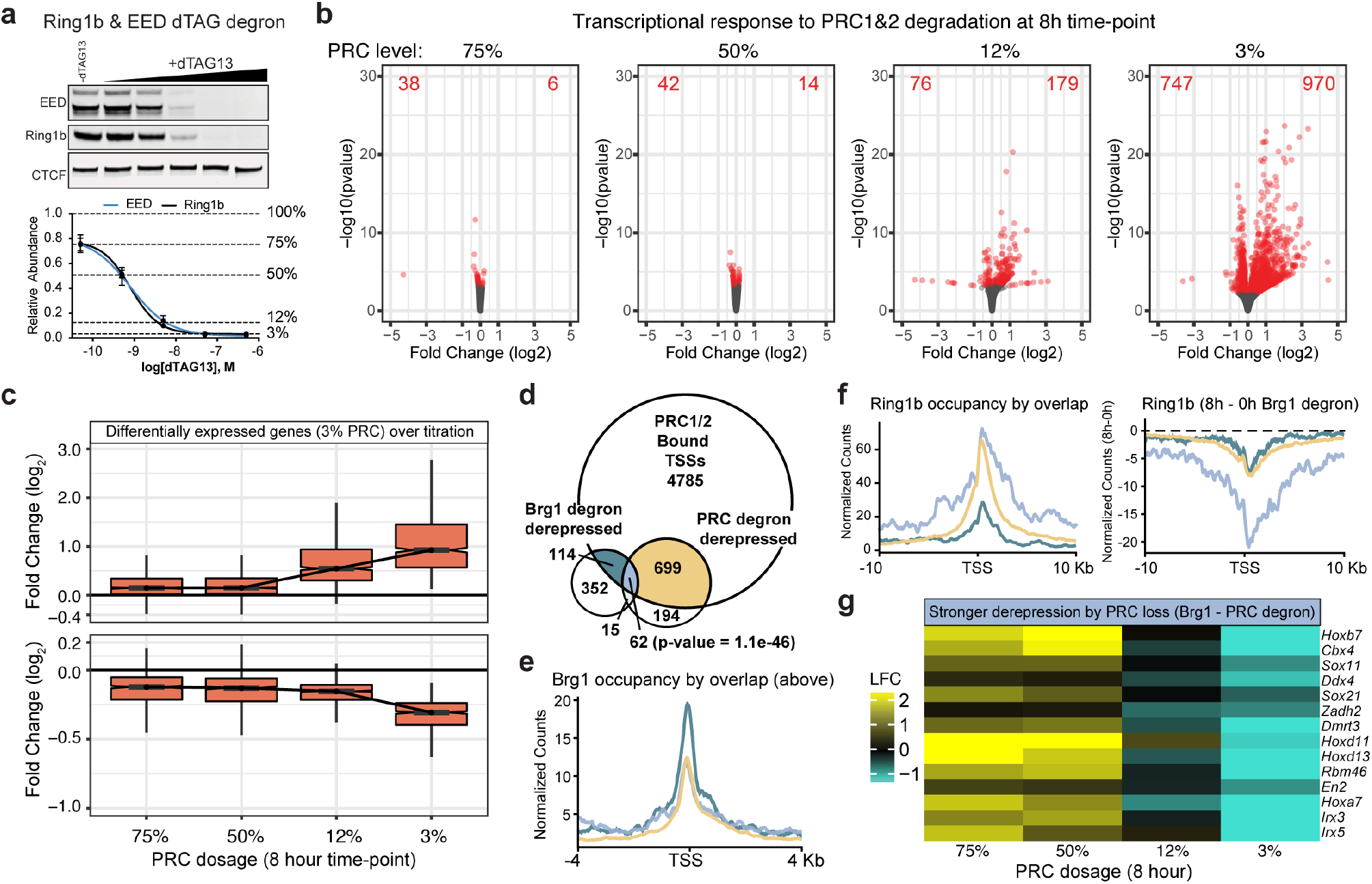
BAF promotes repression directly and by genome-wide polycomb redistribution. (a) Representative western (top) and quantification (bottom) of Ring1b (PRC1) and EED (PRC2) degron lines across 10-fold dilution treatment with dTAG13 PROTAC for 8 hours. Error bars depict mean +/- standard error of the mean from four cell lines. (b) Transcriptional response to degrading PRC1/2 to 75, 50, 12, and 3% of WT levels for 8h. Volcano plots depict gene expression changes across four PRC doses with differentially expressed genes colored red (FDR-corrected P < 0.05) from n = 4 cell lines. (c) Fold change in expression level (Log_2_) for genes that are differentially expressed at 3% PRC dosage (FDR-corrected *P* < 0.05) across all doses (increased n=970, decreased n=747). (d) Significant overlap between genes bound by PRC1/2 (Ring1b and Suz12 +/- 2kb from TSS), Brg1 degron derepressed (8h), and PRC1/2 degron derepressed (3% dosage, 8h). Overlap of PRC-bound, Brg1, and PRC upregulated significance calculated by Fisher’s exact test (p-value = 1.1e-46). (e) Brg1 occupancy is substantially higher over PRC1/2 bound genes solely derepressed in the Brg1 degron compared to genes derepressed by PRC or PRC and Brg1 degradation. (f, left) Higher and broader polycomb domains over genes derepressed by both Brg1 and PRC degron than genes derepressed by Brg1 degron alone. (f, right) Broad domains over genes derepressed by both Brg1 and PRC degron show substantially more redistribution in the Brg1 degron. (g) Heatmap depicting the Log2 fold change in expression between Brg1 degron and each PRC degron dose for select genes more strongly derepressed by PRC than Brg1 degradation. All experiments were conducted in serum/LIF.

We next compared the effects of the Brg1 and PRC1&2 degron at 8h. Our ChIP-seq analysis identified 5,660 genes with both Ring1b and Suz12 at their TSS (+/- 2kb). Of these genes 699 were derepressed by maximum PRC1&2 depletion, 114 by Brg1 degradation, and 62 genes were derepressed by both (Fig. 5d, Fisher’s exact test p-value = 1.1e-46). This confirmed that Brg1 degradation has both divergent and convergent (~35% of Brg1 derepressed genes that are bound by PRC) effects with PRC1&2 depletion. Considering that Brg1 depletion causes rapid transcriptional responses that cannot be explained by downstream feedback mechanisms, we focused on direct mechanisms of repression to explain these results. We found that BAF and polycomb occupancy overlapped extensively but PRC1 deletion had almost no effect on BAF binding (Supplementary Fig. 7) and that BAF was bound to the promoter of all differentially expressed genes, including those with high PRC1 and PRC2 (Supplementary Fig 8e,f). So, we hypothesized that conventional BAF remodeling activity might contribute to repression, in addition to redistributing polycomb (similar to results in Fig 4e). In support of this, Brg1 occupancy was substantially higher at polycomb bound genes that were derepressed by Brg1 alone than genes derepressed by Brg1/PRC1&2 and PRC1&2 depletion (Fig. 5e) (see also Supplementary Fig 8g, for a distinct ChIP approach with a highly similar result). Consistent with these results, genes derepressed by Brg1 depletion alone, and both Brg1/PRC1&2 required BAF to repress accessibility at the promoter (Supplementary Fig. 8h, p-value = 1.1e-9).

We next investigated PRC occupancy at genes derepressed by Brg1 alone, PRC1&2 alone, and both Brg1 and PRC1&2. We found that genes derepressed only by Brg1 loss, had substantially less PRC1 (Fig. 5f) and PRC2 (Supplementary Fig. 8i) compared to those also derepressed by loss of PRC1&2, as expected because they also had substantially higher BAF occupancy. While Brg1 depletion induced polycomb loss at all groups, the genes derepressed by both BAF and PRC1&2 depletion showed substantially greater reduction across a broader range (>20kb). These genes were also modestly dosage sensitive (Supplementary Fig. 8j), where a subset exhibited much stronger derepression by PRC1&2 depletion than BAF, including a few *Hox* genes (Fig. 5g). It’s likely that these genes require greater PRC1&2 depletion than the ~20-60% loss seen by redistribution (Figure 2a,b,g and Supplementary Fig. 3d) because BAF also contributes modestly to their repression. Our results point to a mechanism of BAF-PRC opposition, in which BAF facilitates repression both by directly suppressing accessibility at TSSs and by globally redistributing polycomb across the genome.

### Increased PRC1 dosage inhibits Brg1 degron mediated derepression

To evaluate the contribution of polycomb redistribution, we reasoned that if we conduct the opposite of our PRC1&2 degron experiments and instead overexpress polycomb, then the increased dosage should be sufficient to both redistribute and maintain repression of Hox genes when Brg1 is degraded. This is inherently challenging considering the complexity of the polycomb system. Because it has not been feasible to increase the dosage of each subunit of multisubunit complexes, we sought to overexpress a minimal complex that has a dominant role in repression. To accomplish this, we overexpressed a minimal variant PRC1 complex containing Ring1b and PCGF1, that has the highest H2AK119ub deposition activity, which is essential for polycomb-mediated repression in ESCs^48–50^. Using this strategy, we obtained ~2x increase in H2AK119ub1 levels over the empty vector control, despite ~10-fold RNA overexpression from the strong EF1-a promoter (Fig. 6a). Next, we conducted a series of 8h Brg1 degron experiments to test the effect of vPRC1 overexpression (vPRC1OE) on Brg1-degron mediated derepression. In both vector and vPRC1OE cells, we obtained similar Brg1 degradation efficiencies (95+/- 3.5 and 91+/- 3.0 percent degraded respectively). Yet, overexpressing variant PRC1 significantly inhibited the derepression caused by Brg1 degradation for 10/14 genes that were amenable to qRT-PCR (Fig. 6b, P < 0.05). To further explore the influence of PRC dosage on Brg1 degron mediated derepression, we compared 2i vs. serum conditions at a few genes. Consistent with other reports, 2i culture resulted in a net increase in H2AK119ub (~1.5-fold) and H3K27me3 (~6-fold) (Supplementary Fig. 9a) ^51,52^. Consistent with our overexpression experiments, we found that *Hox* genes were derepressed less efficiently in culturing conditions with increased PRC activity (~3 to 16-fold) (Supplementary Fig. 9b). Thus, increased dosage of just a minimal two-subunit variant PRC1 complex or both PRC1/2 is sufficient to inhibit Brg1 degron mediated derepression, consistent with our model that BAF frees polycomb for passive accumulation across the genome.

**Figure 6:**
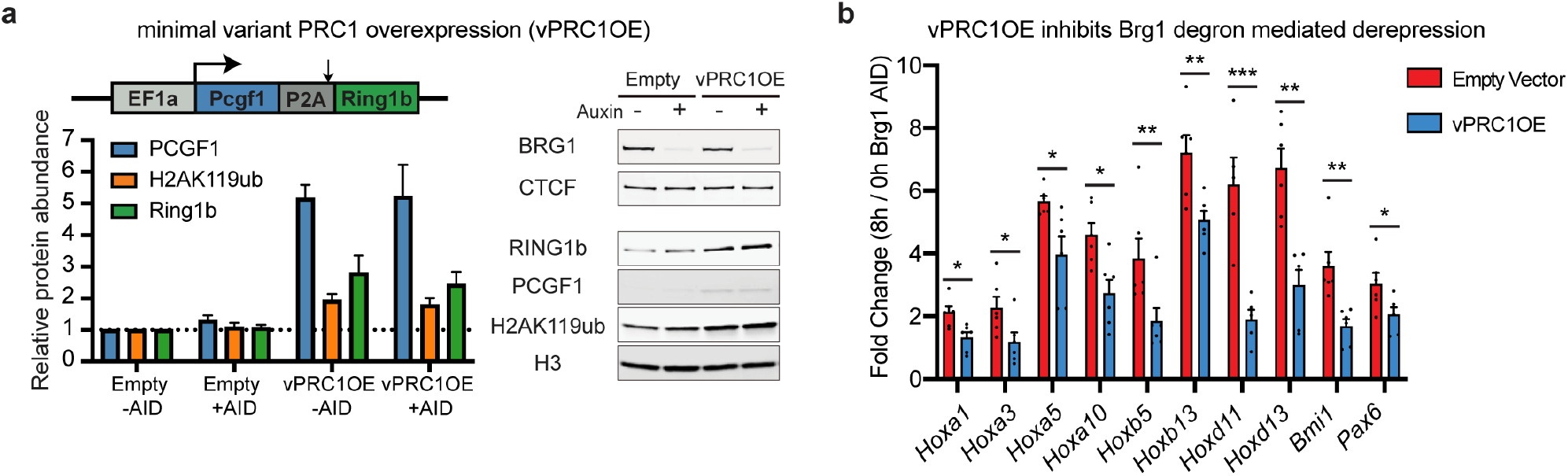
Increased PRC1 dosage inhibits Brg1 degron mediated derepression. (a) Western blot quantification of PCGF1, H2AK119ub, and Ring1b relative to vector only −AID control cells. (right) representative western blot showing Brg1 degron and minimal vPRC1 overexpression (b) Fold change (8h / 0h Brg1 AID) for cells stably expressing either empty vector or variant PRC1. n = 6 biological replicates from two transductions. Significance was calculated by individual Student’s t test (*P ≤ 0.05, **P ≤ 0.01, ***P ≤ 0.001). Error bars represent S.E.M.

## Discussion

Here, we used a targeted degradation approach to comprehensively dissect the BAF-polycomb axis with high temporal precision in mESCs. Our studies indicate that BAF directly promotes polycomb-mediated repression through conventional remodeling activity and by evicting PRC1 and PRC2 such that they can accumulate at distal sites, such as the four hox clusters (Fig. 7). This mechanism of repression reconciles observations that BAF is involved in both polycomb antagonism (reviewed in ref. ^53^) and repression in stem cells and cancer^16–18^, such that direct polycomb eviction allows accumulation over distal sites and hence helps maintain repression of the four hox clusters as well as other developmental genes. It’s not formally possible to completely rule out a direct role for BAF in polycomb loading. Yet, our study revealed that BAF inhibits accessibility where polycomb occupancy is maintained, instead of promoting access, which is inconsistent with BAF acting as a loading factor. Additionally, direct chemically induced BAF recruitment on the time-scale of minutes only leads to polycomb eviction, not loading^11–13^. While, the magnitude of PRC1 and PRC2 loss across these 100+ kilobase scale domains by ChIP-seq is incomplete, the level of depletion was sufficient to physically decompact *HoxA* and *HoxD* loci by ORCA, a completely orthogonal method. These results are consistent, with a previous study that showed heterozygous deletion of Ring1b was sufficient to decompact chromatin at *Hox* clusters^29^, but in this case BAF modulates polycomb dosage, and compaction, at a distance.

**Fig. 7:**
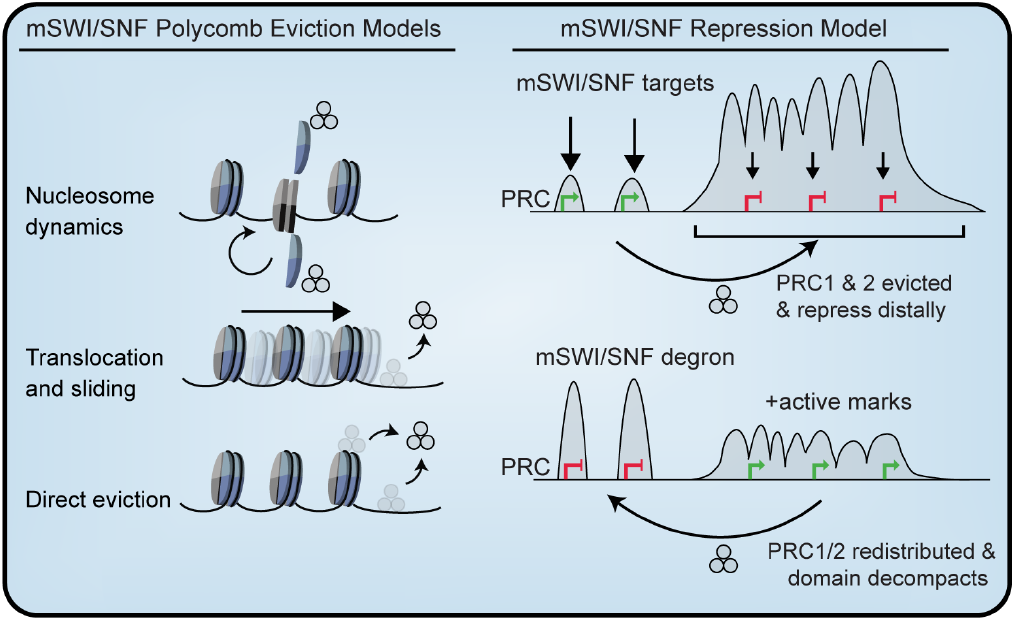
Model and summary of findings. mSWI/SNF polycomb eviction models: a) nucleosome dynamics, b) DNA translocation and nucleosome sliding, c) Direct ATP-dependent eviction. mSWI/SNF Repression Model: mSWI/SNF evicts polycomb, through mechanisms described on the left, enabling accumulation at heavily occupied broad domains like *Hox* clusters, promoting distal repression. Within these repressed domains, BAF is weakly bound at gene promoters supporting another level of repression. When mSWI/SNF is rapidly degraded, PRC complexes accumulate causing redistribution away from heavily occupied sites. This leads to PRC redistribution, physical decompaction of the genome, accumulation of active marks, and derepression.

The fact that we observed derepression of many genes within ~30 minutes of Brg1 degradation indicates that during these experiments, when the dosage of Brg1 reached about 50% of wildtype levels, PRC1&2 began leaving highly-occupied sites and accumulating at sites where Brg1 was no longer evicting them. This result was further supported by short time-course ChIP experiments at the well-studied *Bmi1* locus, showing coincident PRC1&2 depletion and transcriptional derepression. Considering the time needed for genes to be transcribed and spliced, it means that direct PRC1 eviction by the BAF complex must be extremely rapid. This is consistent with studies using CIP to recruit BAF, where PRC1 eviction could be detected in less than 5 minutes, and PRC2 somewhat later^11^. Our studies then call attention to the remarkable and unexpected lability of the PRC-BAF opposition. Thus, our studies support an understanding of BAF-Polycomb opposition that is far more dynamic than the textbook view of rather permanent epigenetic fixation by inherited histone modification.

The BAF-polycomb axis is multifaceted. Both of these groups, of combinatorially assembled complexes, colocalize extensively across the genome^11^ and yet where either is highly bound, the other is mostly excluded. Thus, it’s tempting to conclude that BAF does not function within the most highly occupied polycomb domains, such as *Hox* clusters. Yet, we found that PRC1 deletion had a very minimal effect on BAF binding genome-wide, even within PRC1 domains, revealing that BAF opposes polycomb, but polycomb doesn’t necessarily exclude BAF. Along these lines, our temporally resolved, dose-dependent polycomb degradation approach revealed that there’s more to the repression equation than polycomb redistribution. We found modest BAF binding to promoters of genes derepressed by PRC1&2 degradation and that BAF was required to repress accessibility at the subset of genes derepressed by both Brg1 and PRC1&2. These genes also exhibited disproportionate PRC1 and PRC2 loss upon Brg1 depletion, consistent with a dual mechanism of repression. It’s possible that BAF cooperates with other proteins to facilitate repression such as BRD4S or the NuRD complex, which are known to interact with BAF^54,55^, however future unbiased studies are needed to fully explore this possibility and define all contributors. Nonetheless, the fact that transcriptional derepression is coincident with Brg1 degradation, indicates that secondary effects functioning through altered protein abundance cannot account for the rapid derepression. Furthermore, the polycomb changes induced by Brg1 degradation occurred in the complete absence of transcription and were not restricted to differentially expressed genes. In essence, these results highlight the power of chemical genetics approaches which enabled us to mechanistically dissect a causal role for BAF in polycomb-mediated repression that is lacking in other studies^17,18^.

Polycomb-mediated repression has long been known to be dosage sensitive, such that heterozygous genetic deletion alters expression of *Hox* genes, giving rise to homeotic transformations^4^. Intriguingly, most PRC genes themselves don’t appear to be strongly dosage-sensitive in human disease^56^. An interesting conclusion from our work is that chromatin regulators that antagonize binding, like BAF and possibly others, titrate the effective dosage of all Polycomb complexes. This highlights the importance of gene dosage in regulating the BAF-PRC axis, where even modest overexpression of a single, minimal PRC1 complex was sufficient to inhibit transcriptional derepression caused by Brg1 depletion.

Previously studies showed that BAF evicts PRC1 complexes through a direct interaction with the Brg1 ATPase domain^12^ and that ATPase activity was required for eviction^11,12^. Considering that there are extensive contacts between BAF subunits and the nucleosome core^57^, BAF likely evicts polycomb by three non-mutually exclusive mechanisms: 1) nucleosome dynamics (with polycomb complexes bound to histone tails), 2) DNA translocation and sliding, and 3) direct eviction by BAF (Fig. 7). While, future structural studies are needed to precisely refine the eviction mechanism, there are structural and biochemical data supporting a DNA translocation model for another SWI/SNF subfamily remodeler, Mot1, which pries its substrate TBP off of promoter DNA using ATP hydrolysis^58^. Our studies suggest that this type of antagonism is a determinant of polycomb dynamics in ES cells, which have a hypermobile polycomb fraction^59^. Thus, it’s possible that this dynamic state is especially sensitive to modulation on chromatin, as BAF has been shown to facilitate polycomb-mediated repression in pluripotency, during lineage commitment, and in pediatric brain tumors but not in more differentiated cells.

While individual BAF subunits are highly mutated in specific types of cancer, BAF is almost exclusively mutated in neurodevelopmental disorders but not in other types of human disease^60^. The underlying mechanism for this specificity remains unknown. Our results suggest an explanation by which BAF’s unique dual role in promoting polycomb-mediated repression of genes involved in neurogenesis. A possible explanation for this specificity arises from the observation that the mRNA for certain posterior *Hox* genes such as *Hoxd10-13* increase strongly upon BAF depletion. This would cause the embryo with a mutation in the BAF complex to enter neurogenesis with posterior *hox* genes expressed in developing neural tissue. Based on murine studies this “*hox* confusion” would lead to temporally disordered patterns of gene expression in neural progenitors and abnormal neural development^61^. Consistent with this, Brg1 knockdown in blastocysts was previously shown to result in aberrant *Hox* gene derepression^62^. Recently, Hobert and colleagues found that each class of neurons in C Elegans was found to be delineated by a unique hox code^63^, indicating that confusion of this code would have dramatic effects on neuronal subtypes and their circuitries. While proper examination of this hypothesis would require complex genetic models, it provides a potential explanation for the surprising neural specificity of BAF subunit mutations.

## Online Methods

### Mouse ESC culture

TC1(129) mouse embryonic stem cells were cultured in Knockout™ Dulbeecco’s Modified Eagle’s Medium (Thermo Fisher #10829018) supplemented with 7.5% ES-qualified serum (Applied Stem Cell #ASM-5017), 7.5% Knockout™ Serum replacement (Thermo Fisher #10828-028), 2mM L-glutamine (Gibco #35050061), 10mM HEPES (Gibco #15630080), 100 units mL^-1^ penicillin/streptomycin (Gibco #151401222), 0.1mM non-essential amino acids (Gibco #11140050), 0.1mM beta-mercaptoethanol (Gibco #21985023), and LIF. ES cells were maintained on gamma-irradiated mouse embryonic fibroblast (MEF) feeders for passage or gelatin-coated dishes for assays at 37°C with 5% CO2, seeded at ~3.6×10^4^/cm^2^ every 48 hours, with daily media changes. 2i/LIF conditions were as previously described^64^. To degrade Brg1, osTIR1 was induced overnight with 1.0 μg/mL doxycycline and media containing 0.5 mM 3-indoleacetic acid (Sigma # I2886) and doxycycline was added for indicated time-points. To degrade EED and Ring1b, dTAG-13 was added for 8h at 5×10^-7^-5×10^-11^M. All lines tested negative for mycoplasma.

### CRISPR/Cas9 genome editing

Passage 11 TC1(129) mouse embryonic stem cells were thawed onto MEF coated dishes and passaged once on gelatin coated dishes before transfection. 2M cells were nucleofected (Lonza #VVPH-1001, A-013 program) with 8μg HDR template and 4μg PX459V2.0 (Addgene #62988) containing single guide RNAs (below) or 2μM RNP (IDT #1081058) for dTAG-Ring1b. HDR templates contained 0.5 or 1kB homology arms flanking AID* or FKBP^F36V^ degradation and epitope tags, which were inserted with a flexible (GGGGS)_3_ linker between endogenous protein and tag. Brg1 and EED were tagged at the C-terminus and Ring1b was tagged at the N-terminus. Following transfection 0.5M cells were seeded onto 6cm dishes coated with DR4 MEFS and cultured for 24 hours before puromycin selection (1.0 μg/mL, 1.25 μg/mL, or 1.5 μg/mL) for 48 hours or 1×10^3-5^ cells without selection for RNP. Single colonies were manually picked, dissociated with trypsin and expanded on MEFs. Colonies containing homozygous insertions were confirmed by PCR and western blotting. Guide sequences are as follows: Brg1 sgRNA: TTGGCTGGGACGAGCGCCTC, EED sgRNA: TGATGCCAGCATTTGGCGAT, Ring1b sgRNA: TTTATTCCTAGAAATGTCTC.

### Lentiviral preparation and delivery

HEK293T cells were transfected with gene delivery constructs and packaging plasmids Md2G and psPAX2 using polyethylenimine (PEI). Two days post-transfection the media was collected, filtered, and centrifuged at 50,000 × *g* for 2 hours at 4 °C. The concentrated viral pellet was resuspended in PBS and used directly for transduction or frozen at −80°C. Lentivirus encoding rtTA and TRE-osTIR1 were added to low-passage Brg1-AID* edited cells and selected with 1μg mL^-1^ puromycin and 100μg mL^-1^ hygromycin B.

### RNA isolation and qRT-PCR

Cells were dissociated with trypsin, quenched with media, washed with PBS, and immediately resuspended in Trisure (Bioline # BIO-38033). Total RNA was isolated following manufacturer’s guidelines, digested with DNaseI (Thermo Fischer #18068015), and digestion reaction was cleaned up with acid-phenol:chloroform. cDNA was synthesized from 1μg RNA using the sensifast kit (Bioline #BIO-65054). Primer sequences are in (Supplemental Table 1) and was normalized to GAPDH using the ΔΔCt method.

### RNA-seq and data analysis

RNA sequencing libraries were made from 1μg RNA (RIN > 9) using the SMARTer kit (Takara Bio # 634874) which produces stranded libraries from rRNA depleted total RNA. Libraries were amplified with 12 PCR cycles, quantified with Qubit, and size distribution was determined by Bioanalyzer. Libraries were sequenced single-end with 76 cycles on an Illumina Nextseq or paired-end with 150 cycles and the first three bases were trimmed with cutadapt^65^ before quantification. Transcript abundances were quantified by Kallisto^66^ using the ENSEMBL v96 transcriptomes. Transcript-level abundance estimates from Kallisto and gene-level count matrices were created using Tximport^67^. Differential expression analysis was conducted with DESeq2 version 1.22.2^68^ using default parameters, after prefiltering genes with low counts (rowSums > 10). Differential calls were made by requiring FDR-corrected *P* < 0.05. Log2 fold changes were shrunk with ASHR ^69^ for plotting volcano plots. Boxplot whiskers represent 1.5x interquartile range. Genome browser tracks were generated by aligning trimmed reads to the mm10 genome with HISAT2^70^ with appropriate RNA-strandedness option and deepTools^71^ was used to generate read depth normalized bigwig files, scaled to the mouse genome size (RPGC), for viewing with IGV^72^. Gene ontology analysis was conducted with g:Profiler R client^73^. Gene overlap statistics were calculated using GeneOverlap R package.

### ChIP and ChIP-qPCR

ChIP experiments were performed essentially as described^13^. Briefly, at the end of the time-point 30M cells were fixed with 1% formaldehyde for 12 min, quenched, and sonicated with a Covaris focused ultrasonicator (~200-800bp). Sonicated chromatin was split into separate tubes (~5-10M cell equivalent per IP) and incubated with 5μg primary antibody and 25μL protein G Dynabeads (Life Technologies 10009D) overnight. Beads were washed 4 times and DNA was isolated for qPCR or library preparation. Antibodies are listed in supplemental Table 2. Primer sequences for qPCR experiments are in Supplemental table 1.

### ChIP-seq and data analysis

ChIP-seq libraries were made using NEBnext Ultra II (NEB E7013) with ≤ 12 PCR cycles. Libraries were quantified with Qubit, and size distribution was determined by Bioanalyzer. Libraries were sequenced single-end with 76 cycles on an Illumina Nextseq. Reads were aligned to the mm10 genome with Bowtie 2^74^ version 2.3.4.1 with the --very-sensitive option. Alignments were removed with the following criteria: quality score < 20, PCR duplicates, secondary alignments, and supplemental alignments. Peaks were called with MACS2 using default parameters and requiring that q < 0.01. Peaks from control and treated datasets within +/- 1kb were merged, filtered against the mouse blacklist^75^ and the number of reads overlapping these peak sets were compared for differential peak calling. Differential peak calls were determined using DESeq2 after pre-filtering peaks with low counts (rowSums > 400 for Ring1b & Suz12 with four biological replicates, and rowSums > 100 for histone marks with two biological replicates. Differential calls were made by requiring FDR-corrected *P* < 0.1. For browser snapshots, replicate BAM files were merged and read depth normalized genome coverage files (bigwig) were generated with deepTools^71^, scaled to the mouse genome size (RPGC). For subtractions, normalized bigwig files were subtracted using deepTools^71^ (8 hour – 0 hour) for visualization by heatmap, metagene plot, or by browser (IGV).

### RNA polymerase inhibition

Transcription initiation was blocked by treating cells with 10 μM triptolide (Sellec S3604). To validate triptolide treatment, quantitative RT-PCR was performed by comparing derepressed genes in triptolide, triptolide and IAA, to IAA only cells, normalized to U6 snRNA using the ΔΔCt method.

### ORCA Imaging and data analysis

The primary probes tiling the *HoxA* and *HoxD* DNA regions at 5kb resolution (Supplemental data), were designed as previously described^32^. Probes were amplified from the oligopool (CustomArray) and amplified according to the protocol described in^28,32^.

In preparation for imaging, mESC cells were plated on 40-mm glass coverslips (Bioptechs) coated with 0.1-0.2% Gelatin, and fixed on the following day in 4% PFA in 1xPBS for 10 min. The hybridization and imaging were performed as previously described (Mateo et al. 2019). Briefly, for primary probe hybridization, cells were permeabilized for 10min with 0.5% Triton-X in 1xPBS, the DNA was then denatured by treatment with 0.1M HCL for 5min. 2ug of primary probes in hybridization solution was then added directly on to cells, placed on a heat block for 90C for 3min and incubated overnight at 42C in a humidified chamber. Prior to imaging, the samples were post-fixed for 1h in 8%PFA +2% glutaraldehyde (GA) in 1xPBS. The samples were then washed in 2xSSC and either imaged directly or stored for up to a week in 4C prior to imaging. For imaging samples were mounted into a Bioptechs flow chamber, and secondary probe hybridization and step by step imaging of individual barcodes, and image processing was performed as in^32^.

Image analysis was performed as described in^32^. For all analysis in Fig. 3, we excluded all barcodes that had a low labeling efficiency in either control or AID treated condition (labelled in less than 10% of the cells). This resulted in a final of 52 barcodes for *HoxA* and 28 barcodes for *HoxD*. For comparison with HiC, mESC data from^33^ was downloaded from Juicebox^76,77^ and matched to ORCA barcode coordinates by finding the closest genomic bins in HiC, and removing bins corresponding to excluded barcodes. ORCA data was processed to calculate contact frequency across all cells (Supplemental Fig. 4 A, B), where contact frequency was computed by calculating the fraction of cells where the probes were within a 200 nm distance. We use ‘cell’ to refer to all detected spots, with ~2 spots per cell, corresponding to each allele. For calculation of median probe distance (to generate Fig. s 4E, F) we calculated the median distance across all cells in each condition. For both *HoxA* and *HoxD* imaging, we obtained more cells in the control condition (1238 for *HoxA*, 2419 for *HoxD*) than in the AID treated condition (587 for *HoxA* and 1183 for *HoxD*). To control for sample size, we randomly split the cells in the control dataset into 2, calculated median of pairwise probe distances for each subset and computed the difference in pairwise distances between the two halves and between each control subset and Brg1 depleted cells (Supplemental Fig. 3D, E).

## Data availability

All sequencing data have been deposited in GEO (accession number GSE145016)

## Acknowledgements

We thank members of the Crabtree lab for insightful comments and discussions over the course of the study. We are grateful to Andrey Krokhotin, Srinivas Ramachandran (CU Anschutz), and Vijay Ramani (UCSF) for providing critical comments on the manuscript and for helpful discussions. This study was supported by NIH grants R01CA163915 (G.R.C), R37NS046789 (G.R.C), DP2GM132935 (A.N.B.), the Howard Hughes Medical Institute (G.R.C), Swiss National Science Foundation (SNSF) postdoctoral fellowship (S.M.G.B.), the Walter V. and Idun Berry Postdoctoral fellowship program (C.M.W. and T.A.), and the Sir James Black postdoctoral fellowship (C.M.W. and S.M.G.B.).

## Author Contributions

C.M.W conceived of the project, did experiments, and analysis. T.A. performed ORCA experiments and analysis. S.M.G.B., J.K.G, and B.Z.S. performed experiments. G.R.C and A.N.B. designed experiments and supervised the project. C.M.W wrote the paper with assistance from all authors.

## Supplementary Figures

**Supp. Fig 1.**
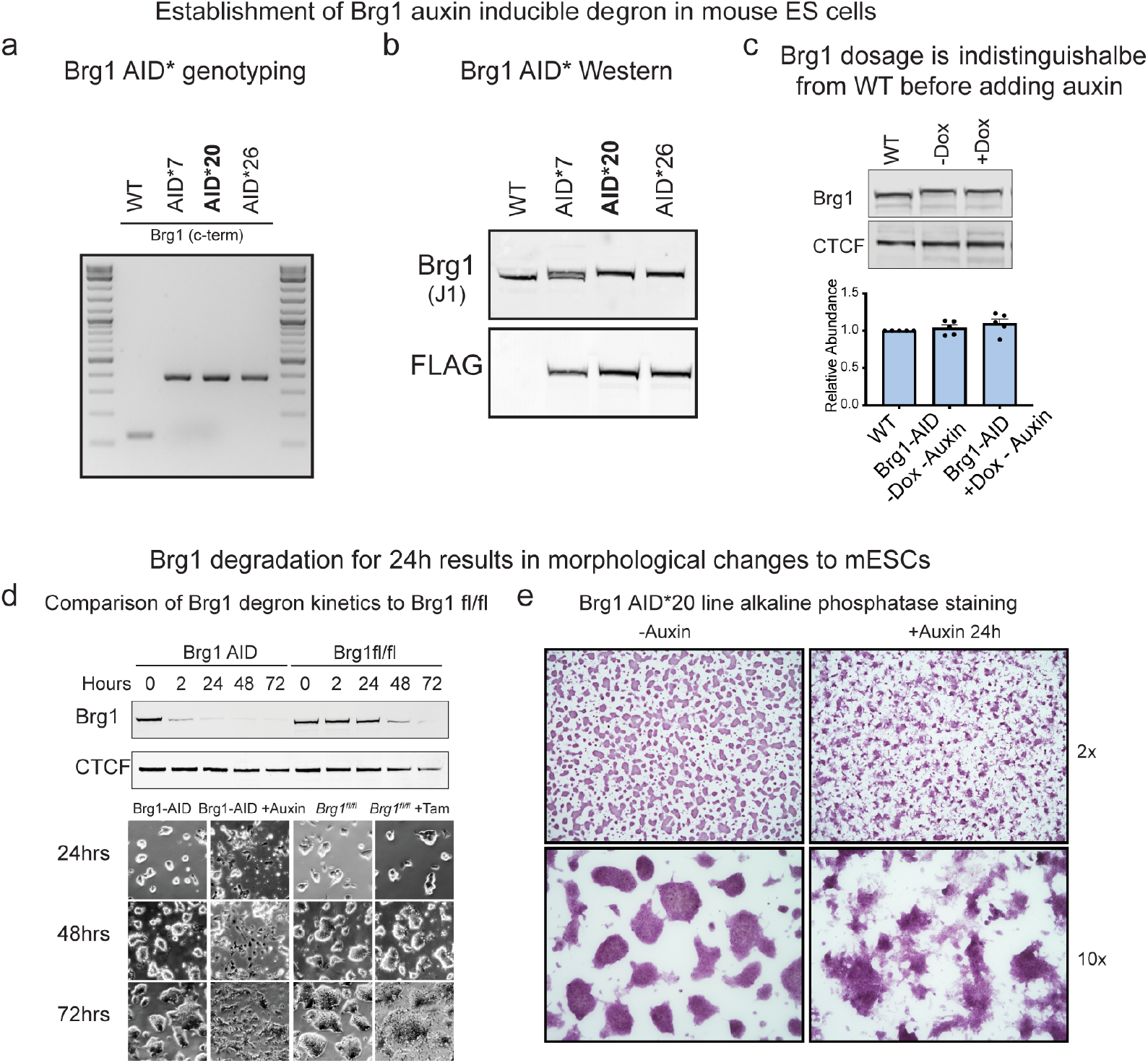
Extended characterization of Brg1 AID degron, related to Fig. 1. (a) Genotyping PCR for three homozygous knock-in clones (AID*20 was used for most experiments in the manuscript). (b) Western blot of wholecell extracts from knock-in lines showing 8.8kDa shift in migration from c-terminal tag (G4S)3-AID*-G4S-3xFLAG. Line 20 and 26 have homozygous insertions in frame that were also confirmed by Sanger sequencing. (c) Representative western blot and quantitation of Brg1 abundance in parent line compared to Brg1-AID* +/- osTIR1 induction (-auxin). Error bars represent mean +/- standard deviation from five biological replicates. (d) Comparison of Brg1 depletion kinetics between AID and Cre-lox genetic deletion by western blot and changes to colony morphology that occur at the 24-hour time-point. Cells were only grown on the same dish for 72h to illustrate timing for Brg1^fl/fl^ colony morphology changes. (e) Alkaline phosphatase staining of Brg1 AID* 20 line on gelatin coated dishes, showing that changes to colony morphology occur ~24h.

**Supp. Fig 2.**
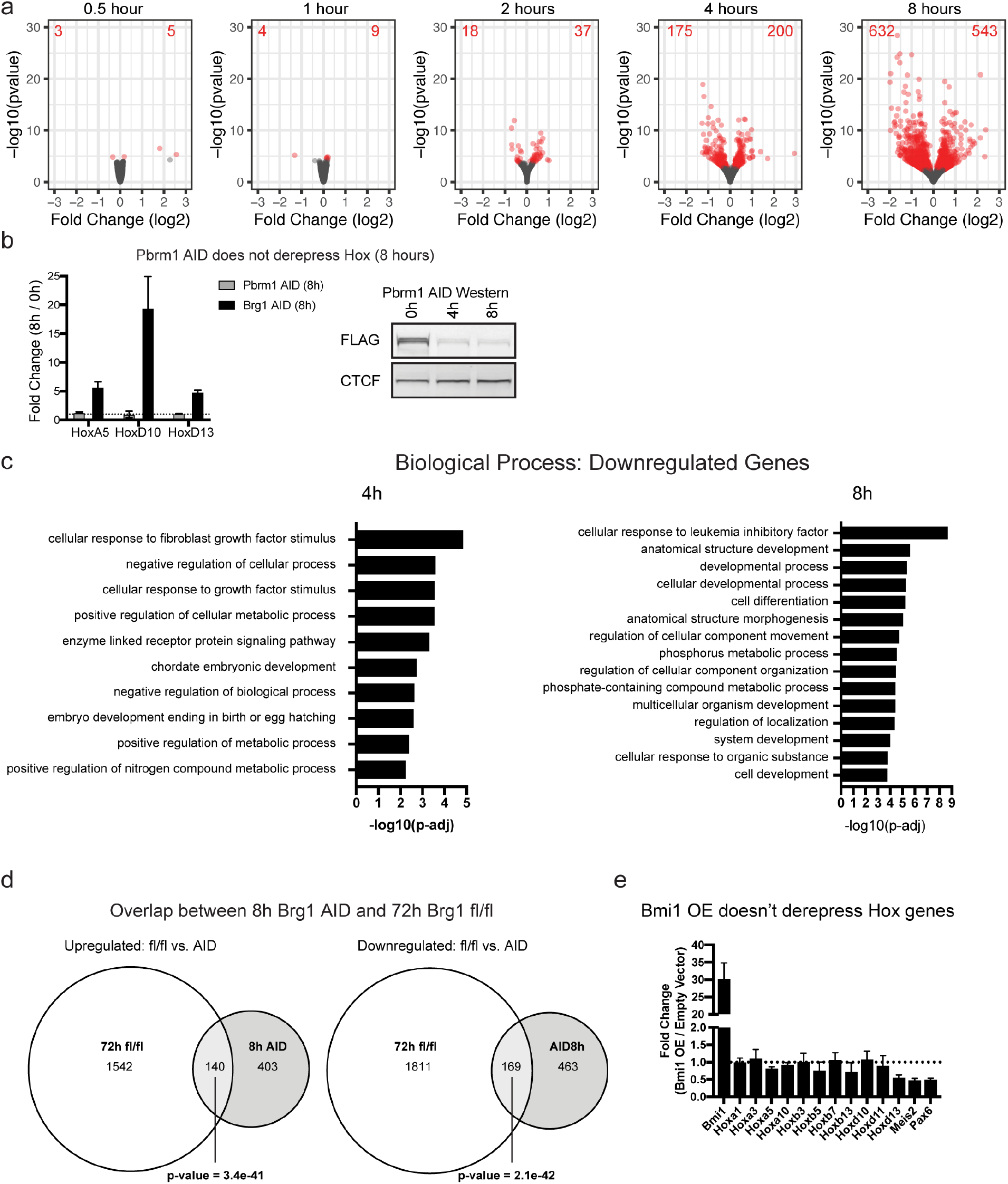
Extended summary of RNA-seq results, related to Fig. 1. (a) Volcano plots depicting gene expression changes at 0.5, 1, 2, 4, and 8 hours of Brg1 degradation. Differentially expressed genes are colored red (FDR-corrected P < 0.05) from four biological replicates for each time-point. (b) Pbrm1 degradation with auxin, does not derepress *Hox* genes, revealing that pBAF is not required for their repression and controls for auxin, Tir1 expression, and proteasomal degradation in the Brg1 degron. Error bars are mean +/- standard error for n=3 replicates. (c) Gene set enrichment analysis for differentially downregulated genes at 4 and 8-hour timepoints. (d) Overlap of differentially expressed genes in 72h Brg1 fl/fl from^12^ and 8h Brg1 AID calculated by Fisher’s exact test (p-value = 1.1e-46). (e) Strong lentiviral Bmi1 overexpression doesn’t derepress *Hox* genes or other genes derepressed by Brg1 degron.

**Supp. Fig 3.**
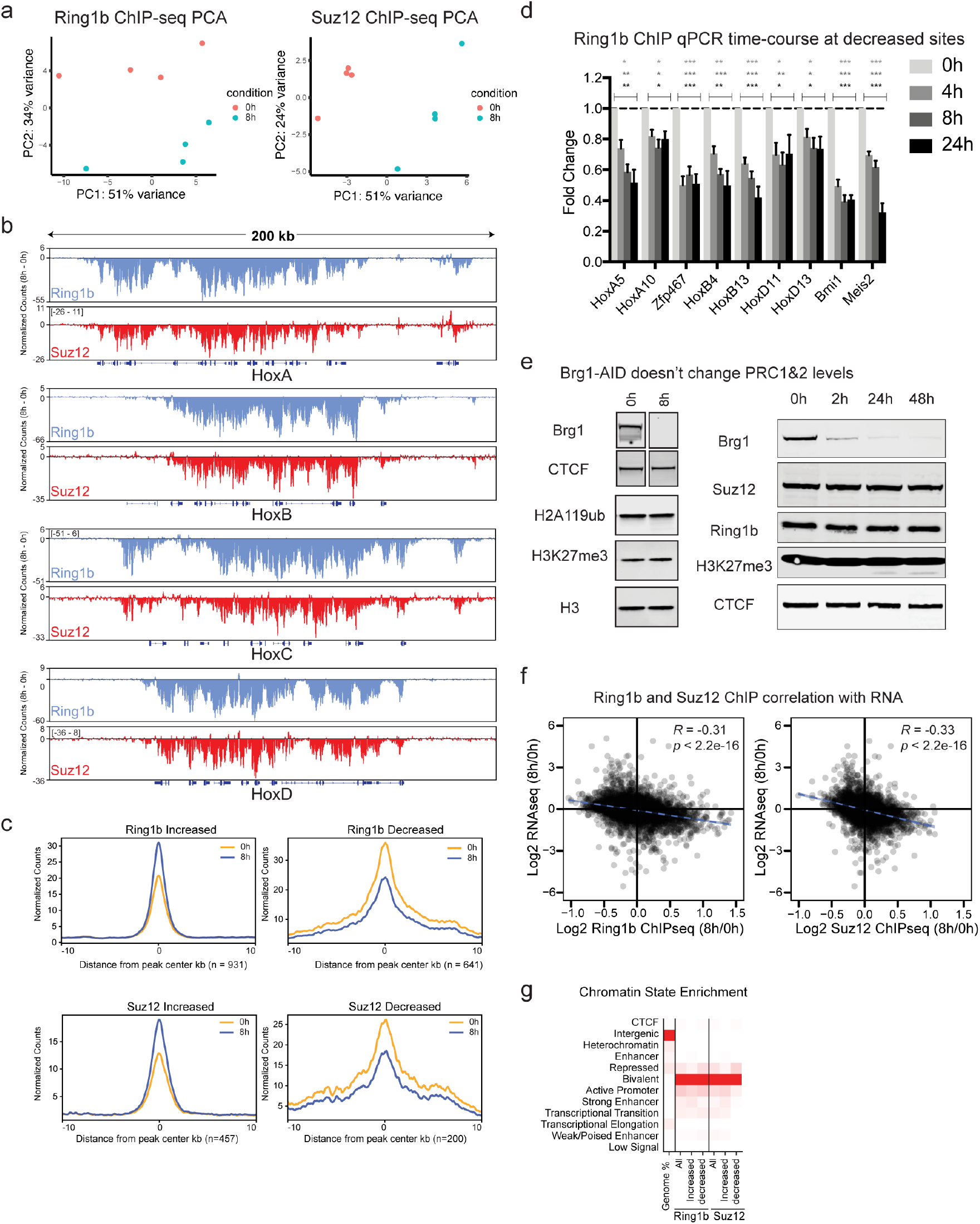
Brg1 degradation results in quick redistribution of PRC1&2 but doesn’t affect net dosage, related to Fig. 2. (a) Principal component analysis of Ring1b and Suz12 ChIP-seq (n = 4 replicates) (b) Browser snapshot of the difference in normalized counts (8h – 0h) for Ring1b and Suz12 at *Hox A, B, C*, and *D* depicting net loss across all four clusters. (c) Average metagene plots of normalized counts (to read-depth) at sites that were significantly increased or decreased by DESeq2 analysis (FDR-corrected P < 0.1) and normalized to peak intensity, providing additional confirmation that normalization is correct. (d) ChIP qPCR timecourse at 4, 8, and 24h Brg1 degradation for Ring1b at sites that were significantly decreased by ChIP-seq. Error bars depict standard error of the mean from four biological replicates. (* = p<0.05, ** = p<0.005, *** = p<0.0005; t test with Holm-Sidak multiple comparison correction). (e) (left) Representative western blot showing Brg1 degradation efficiency and that H2AK119ub and H3K27me3 marks don’t change at 8h time-point (ChIP-seq time-point for the complexes that deposit these marks). (right) Representative western showing Brg1 degradation efficiency over a longer timecourse and that Suz12, Ring1b, and H3K27me3 marks don’t noticeably change even at 48h. (f) Correlation of all Ring1b and Suz12 peaks that are +/- 2kb from gene transcription start sites and expression changes at 8h Brg1 degradation. Correlation and p-values were obtained from Pearson’s product moment correlation. (g) chromatin state enrichment for all, increased, and decreased peak changes for Ring1b and Suz12.

**Supp. Fig 4.**
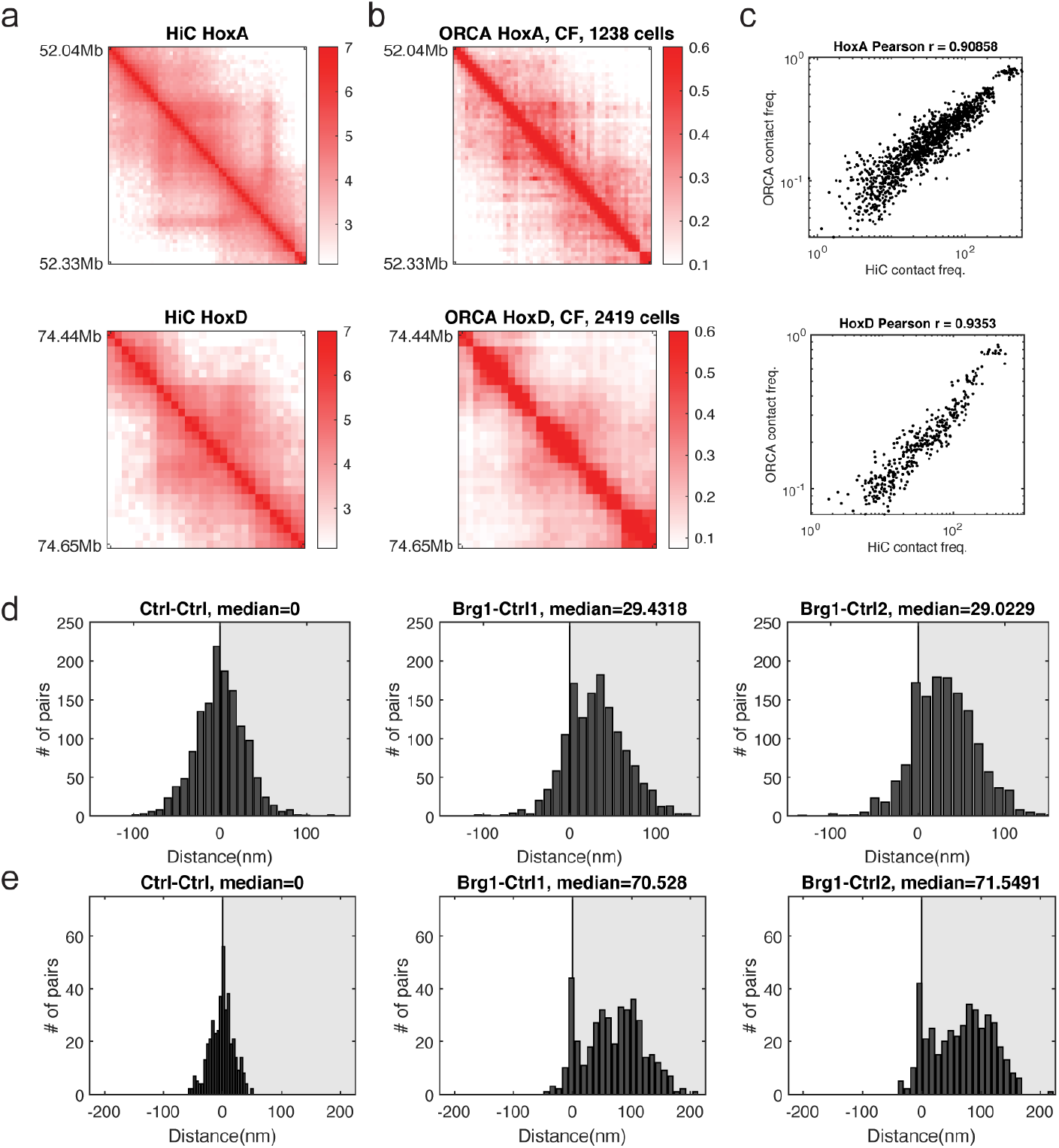
Comparison of ORCA to Hi-C, related to Fig. 3. (a) Average contact frequency as measured by high resolution Hi-C from 2i grown cells ^33^. (b) Average contact frequency as measured by ORCA (150-nm threshold). (c) Pearson’s correlation and Pearson’s r were computed using all unique pairwise combinations of X,Y barcodes measured. (d) Histograms depicting the difference in median distance for barcode pair combinations at *HoxA*. (e) Histograms depicting the difference in median distance for barcode pair combinations at *HoxD*.

**Supp. Fig 5.**
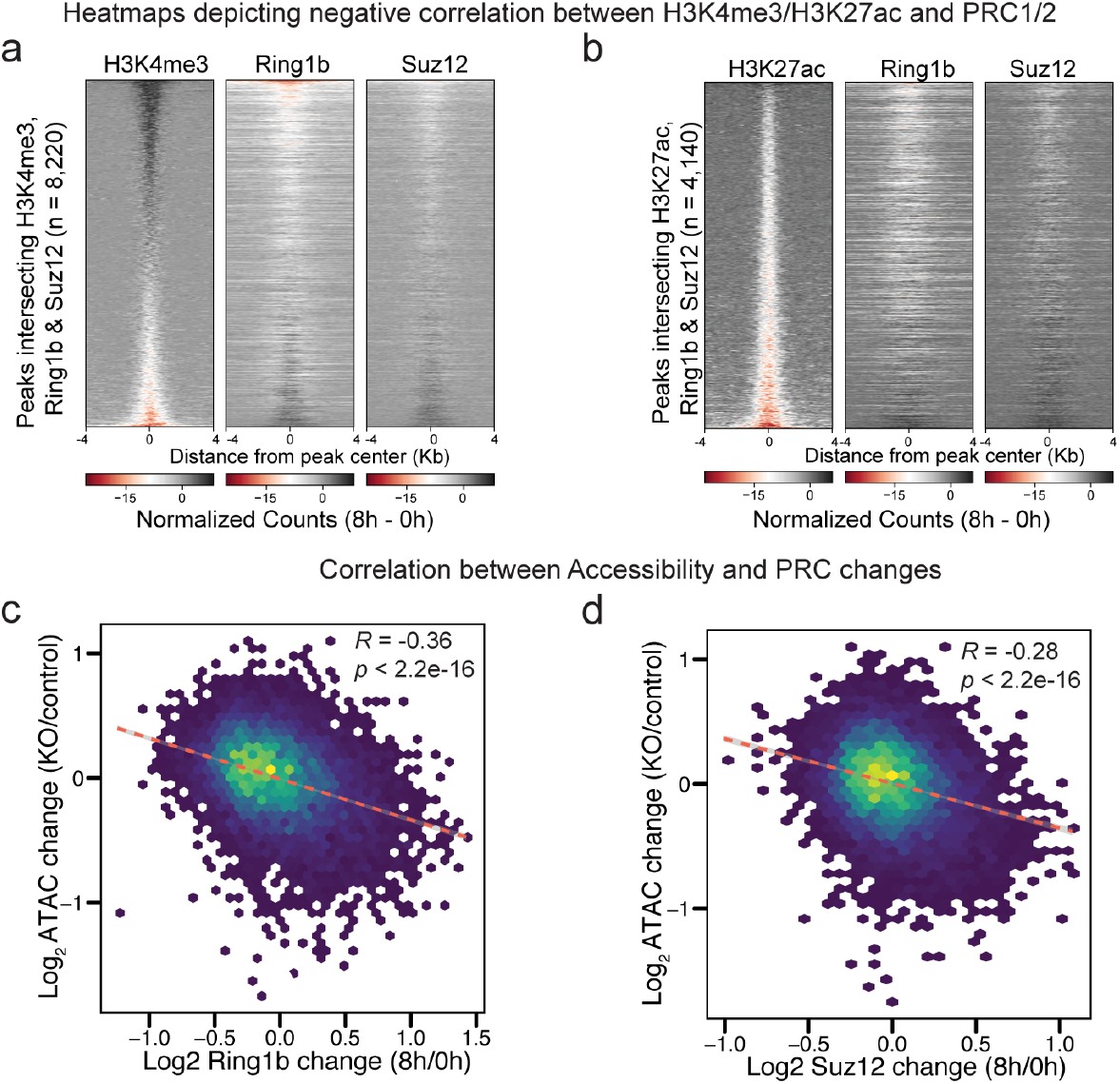
Extended analysis related to Fig. 4. (a) Heatmap depicting the negative correlation between H3K4me3 and Ring1b and Suz12 as the difference in normalized counts (8h – 0h) at peaks (8,220) that intersect H3K4me3, Ring1b, and Suz12 at merged peak center, sorted by decreasing H3K4me3 signal. (b) Heatmap depicting the negative correlation between H3K27ac and PRC1&2 as the difference in normalized counts (8h – 0h) at peaks (4,140) that intersect H3K27ac, Ring1b, and Suz12 at merged peak center, sorted by decreasing H3K27ac signal. (c,d) Scatterplots showing negative correlation between ATAC-seq changes (KO/control) ^41^ and Ring1b (8h/0h) or Suz12 (8h/0h).

**Supp. Fig 6.**
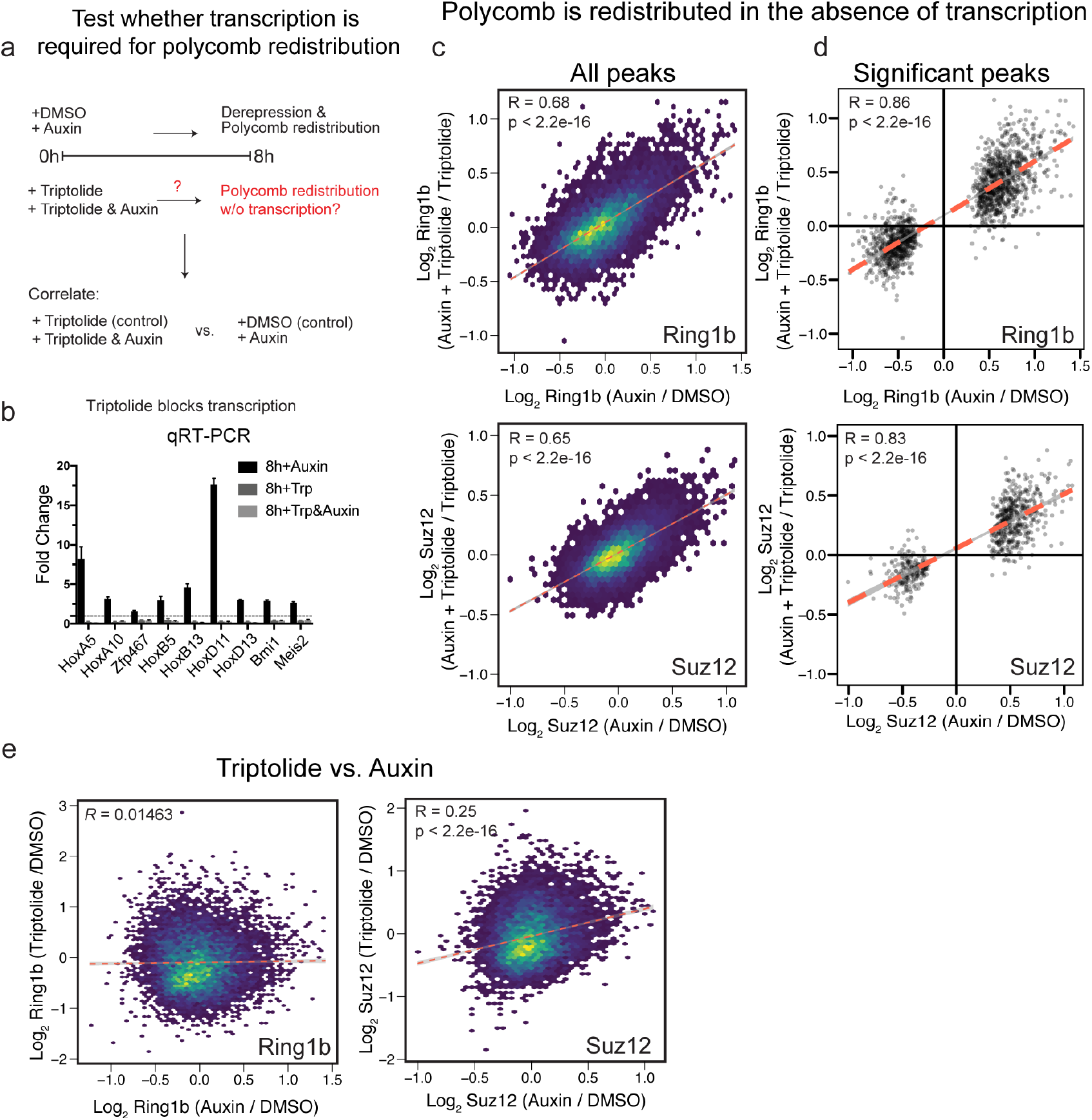
Transcription is not required for polycomb redistribution. (a) Experimental design to test whether polycomb redistribution requires transcription. (b) qRT-PCR depicting fold change by ΔΔCt method with indicated treatments. Scatterplot showing that Ring1b and Suz12 changes following Brg1 degradation are highly correlated despite global inhibition of transcription for all peaks (c) and peaks that were significantly changed by Brg1 degradation alone (d) (FDR < 0.1). (e) Scatterplot of changes to Ring1b and Suz12 upon treatment with auxin (auxin / control) or triptolide (triptolide / control) showing distinct effects from the two treatments. Correlation and p-values were obtained from Pearson’s product moment correlation for c-e, from n = 4 biological replicates at the 8h timepoint.

**Supp. Fig 7.**
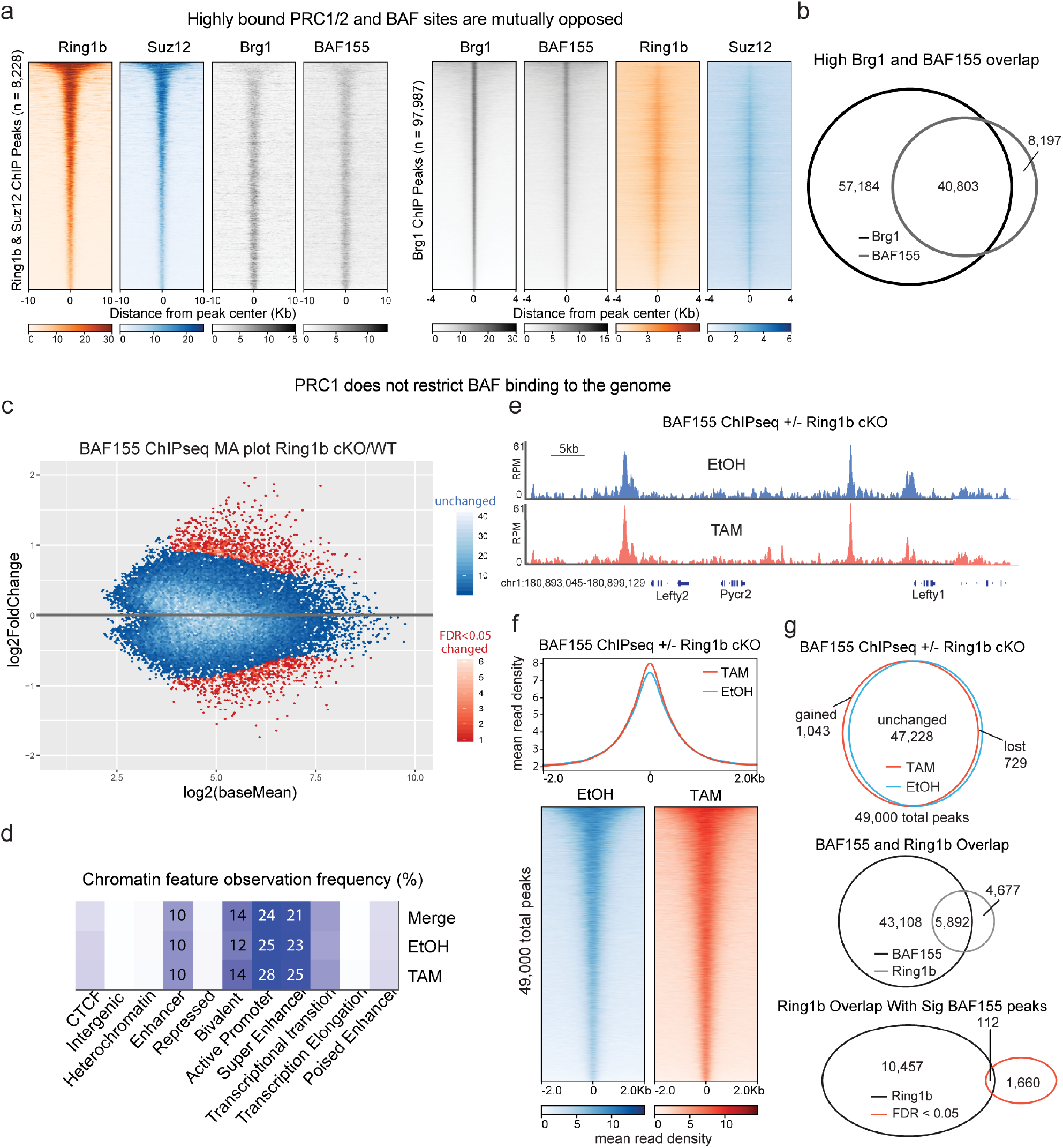
PRC1 does not restrict BAF binding despite extensive overlap. (a) Heatmaps depicting Ring1b, Suz12, BAF155, and Brg1 ChIPseq signal from ^78^ at peaks (8,228) that overlap Ring1b and Suz12, sorted by decreasing Ring1b signal (left) or Brg1 peaks (97,987) sorted by decreasing Brg1 signal (right). (b) High overlap between Brg1 and BAF155 despite different number of peaks called due to antibody quality and read depth. (c) BAF155 ChIP-seq in Ring1b^fl/fl^ x Ring1a^-/-^ x ActinCreER ESCs treated with Tamoxifen or EtOH for 72h. MA plot (TAM/EtOH) highlights large number of unchanged peaks (>95% in blue) (n=2). (d) Analysis of the chromatin features that characterize BAF155 binding sites reveal that Ring1a/b double KO does not alter BAF complex binding, as the complex remains mainly bound to enhancers and bivalent promoters in both control and mutant ESCs. (e) Representative genome tracks showing similar BAF155 levels across the Lefty1/2 locus in Tamoxifen and EtOH treated cells. (f) Metagene plot and heatmaps showing that BAF155 ChIPseq signal is minimally affected by Ring1b deletion (g) Characterization of BAF155 peaks in EtOH and Tamoxifen conditions. Of the 49,000 detected BAF155 peaks in this dataset, less than 5% were significantly changed upon Ring1a/b double KO in ESCs and the small number of gained and decreased sites was similar. Despite 56% of Ring1b peaks overlapping with a BAF peak, only 1% of differentially bound peaks are within a PRC1 domain (bottom).

**Supp. Fig 8.**
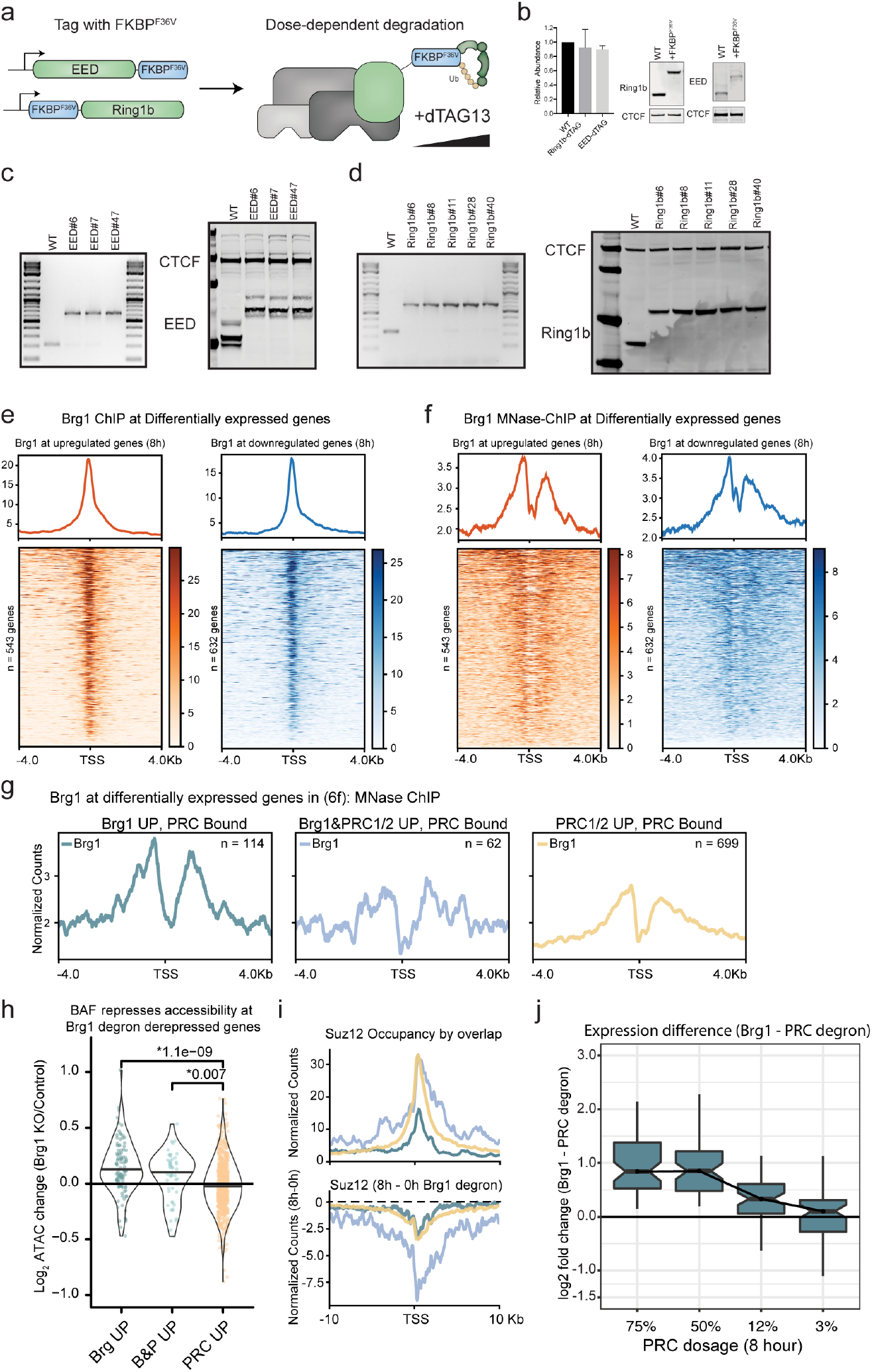
Characterization of EED/Ring1b degron and extended analysis related to Fig. 5. (a) Schematic depicting EED and Ring1b dTAG targeted protein degradation strategy in mESCs, where EED is tagged at the C-terminus and Ring1b is tagged at the N-terminus with FKBP^F36V^ to enable targeted degradation with dTAG13 PROTAC. (b) Representative western blot and quantitation of EED and Ring1b abundance in parent line compared to lines tagged with FKBP^F36V^ Error bars depict mean +/- standard error of the mean from three tagged EED lines and four tagged Ring1b lines. (c) Genotyping PCR and western from whole cell extract showing 14.6kDa shift in migration from C-terminus (G4S)3-FKBP^F36V^-G4S-V5 tag on three different EED clones used for experiments. (d) Genotyping PCR and western from whole cell extract with 14.2kDa shift in migration from N-terminus FKBP^F36V^-G4S-HA-(G4S)3. (e) Brg1 ChIPseq showing localization to the TSS of differentially expressed genes, ChIP from ^78^. (f) Brg1 MNase ChIPseq showing localization to the TSS of differentially expressed genes, ChIP from ^79^. (g) Brg1 MNase ChIPseq showing higher occupancy at PRC bound genes repressed by Brg1 alone. Compare to Fig. 5f. (h) Brg1 represses accessibility at PRC bound genes derepressed by Brg1 degron and Brg1/PRC degron. Accessibility change (Brg1 KO / control) by overlap (Fig. 6f) for peaks +/- 2kb from gene promoter, ATAC data from ^41^ with t-test p-value above. (i,top) Higher and broader polycomb domains over genes derepressed by both Brg1 and PRC degron than genes derepressed by Brg1 degron alone, with Suz12 occupancy shown here. (i, bottom) Broad domains over genes derepressed by both Brg1 and PRC degron show substantially more redistribution in the Brg1 degron. Compare to Ring1b signal in Fig. 6. (j) Expression difference (Brg1 – PRC Log_2_ Fold Change) for genes derepressed by Brg1 and PRC degron, over dosage titration (Most dosage sensitive genes are shown in Fig. 6g).

**Supp. Fig 9.**
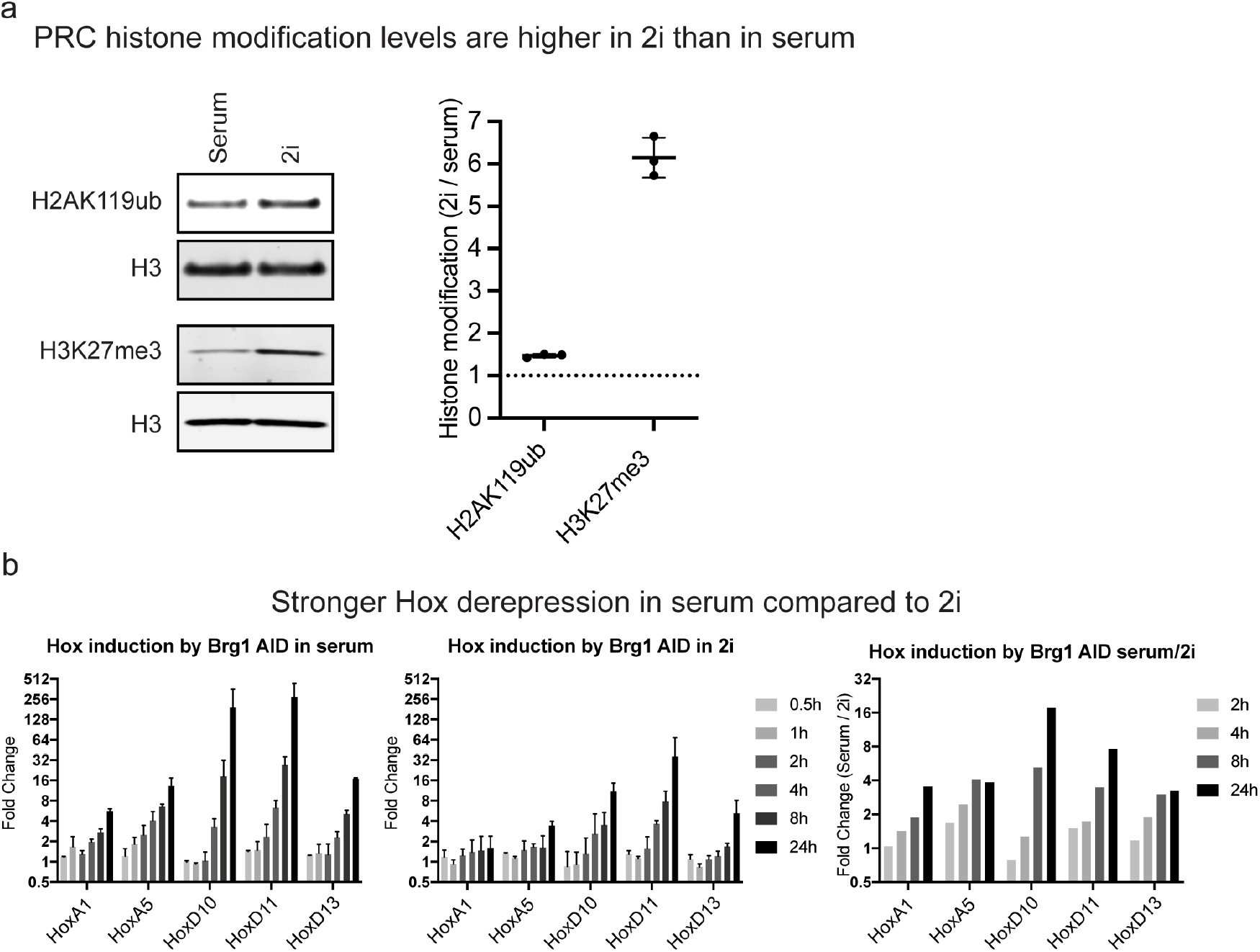
Histone modifications deposited by PRC1 and PRC2 are higher in 2i growth conditions and results in Weaker derepression, related to figure 2. (a) Western blot and relative quantification of H2AK119ub and H3K27me3 normalized to H3 in cells grown in Serum/LIF or 2i growth conditions. (b) *Hox* gene derepression measured by qRT-PCR for cells grown in serum, 2i, and the ratio of serum/2i. Error bars are mean+/- standard deviation from two biological replicates.

